# The role of host phenology for parasite transmission

**DOI:** 10.1101/855031

**Authors:** Hannelore MacDonald, Erol Akçay, Dustin Brisson

## Abstract

Phenology is a fundamental determinant of species distributions, abundances, and interactions. In host-parasite interactions, host phenology can affect parasite fitness due to the temporal constraints it imposes on host contact rates. However, it remains unclear how parasite transmission is shaped by the wide range of phenological patterns observed in nature. We develop a mathematical model of the Lyme disease system to study the consequences of differential tick developmental-stage phenology for the transmission of *B. burgdorferi*. Incorporating seasonal tick activity can increase *B. burgdorferi* fitness compared to continuous tick activity but can also prevent transmission completely. *B. burgdorferi* fitness is greatest when the activity period of the infectious nymphal stage slightly precedes the larval activity period. Surprisingly, *B. burgdorferi* is eradicated if the larval activity period begins long after the end of nymphal activity due to a feedback with mouse population dynamics. These results highlight the importance of phenology, a common driver of species interactions, for the fitness of a parasite.

## Introduction

Behaviors or traits that vary seasonally, termed phenology in the ecological literature, impact both the type and strength of ecological interactions within populations and communities (Miller-Rushing et al. 2010; Bewick et al. 2016; Paull and Johnson 2014; Barber et al. 2016; Burkett-Cadena et al. 2011). For example, seasonal matching between flowering times and pollinator activity periods is a key driver of short- and long-term population dynamics of both plants and insects (Cleland et al. 2007; Gaku et al. 2004; Inouye 2008; Kudo and Ida 2013; Memmott et al. 2007; Hegland et al. 2009). Differences in the seasonal activities of interacting species over time or geography, caused by changes in climatic and environmental features, can result in population extinctions and in population explosions (Cahill et al. 2013; Johnson et al. 2010; Washburn and Cornell 1981; Powell and Bentz 2009; Jepsen et al. 2009; van Asch and Visser 2007; Jepsen et al. 2008). Although the majority of studies focus on the phenology of plants and their interacting species, the seasonal activity of hosts or disease vectors is also likely to have large impacts on the population dynamics of infectious microbes.

The impact of phenology on disease transmission dynamics can be prominent in disease systems involving multiple host species or life-stages because the seasonal match or mismatch of activities between species or stages will determine the frequency and type of pathogen transmission. For instance, consider the cestode *Schistocephalus solidus* that infects young three-spined stickleback fish as an intermediate host, multiplies within the fish before the fish is eaten by the definitive bird host (belted kingfisher) (Clarke 1954; Heins et al. 2016). The parasite reproduces sexually within the bird who defecate parasite eggs that infect juvenile fish (Clarke 1954). This disease system occurs in North American lakes that freeze over winter, causing both fish reproduction and bird migration to be temporally restricted within each year. A temporal mismatch in the bird and fish phenologies, such as fish reproduction occurring prior to the return migration of birds, could therefore reduce or eliminate cestode transmission among its hosts. Further, variation in the environmental cues affecting the seasonal activity patterns of the birds and fish either among lakes or across years is likely to impact disease transmission dynamics. These types of seasonal dynamics are expected to impact parasite fitness in many disease systems, yet the quantitative and qualitative impact of phenology remains relatively under explored (Barber et al. 2016).

Human diseases caused by zoonotic pathogens, those that complete their natural life cycle in wildlife but can infect humans, are likely impacted by the phenology of their wildlife hosts or vectors. Parasites transmitted by hard bodied ticks (family *Ixodidae*) represent a practical case study to examine the impact of phenology on disease systems. The public health importance of diseases transmitted by these ticks, such as Lyme disease, has resulted in expansive field datasets that provide baseline expectations for the transmission consequences of tick phenological patterns, making this a good system to study the effects of the general conceptual issue of how vector phenology drives parasite transmission (Randolph 1999b; Randolph et al. 2000; Ogden et al. 2018). Ixodid ticks have three distinct developmental stages. Larvae, the first developmental stage, hatch uninfected but can acquire *Borrelia burgdorferi*, the etiological agent of Lyme disease, while feeding on an infected host (Fig. 1). Fed larvae molt to nymphs that can then transmit *B. burgdorferi* to small vertebrate hosts (primarily mice, chipmunks, and shrews) during nymphal feeding. Fed nymphs molt to adults that feed on large vertebrates before laying eggs that hatch as larvae. In the Northeastern US, the nymphal stage is active in early summer while larvae from a different cohort feed in late summer, providing an opportunity for *B. burgdorferi* transmission from nymphs to larvae through the vertebrate hosts (Wilson and Spielman 1985). This sequential feeding pattern has been alleged to contribute to higher infection prevalence found in the Northeastern US relative to Southern or Midwestern US, where the sequential activity patterns are less pronounced (Ogden et al. 2018; Brinkerhoff et al. 2011).

**Figure 1:**
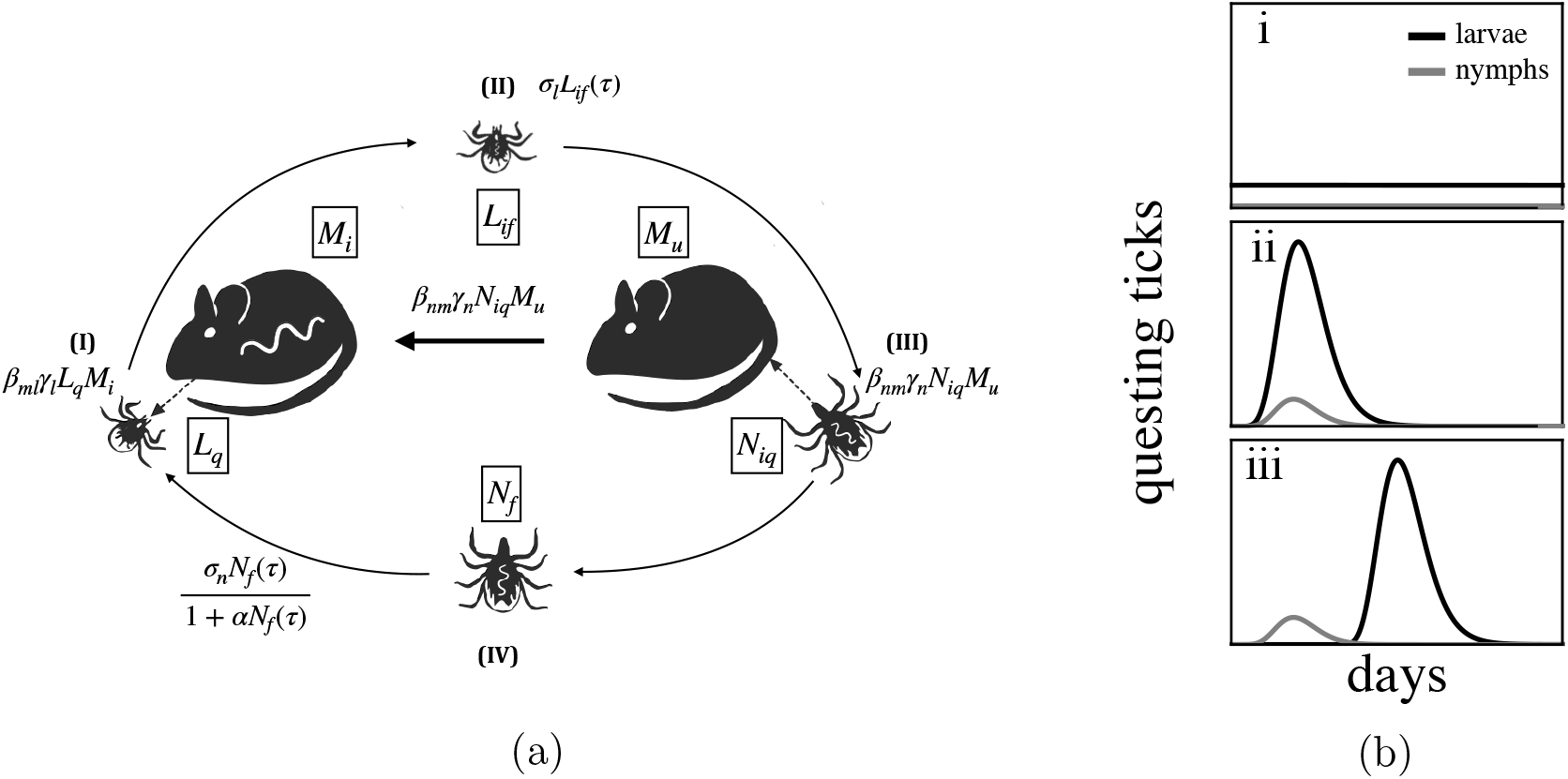
How does tick life-stage phenology impact the transmission of *B. burgdorferi*? (*a*.) Larval ticks hatch uninfected (Piesman et al. 1986; Patrican 1997) and can acquire *B. burgdorferi* by feeding on an infected small animal (*I*). Fed infected larvae molt to nymphs and become active the following year (*II*). Small animals are infected when fed upon by an infected nymph (*III*). Fed nymphs molt to adults and feed on larger animals prior to laying eggs that hatch the following year (*IV*). Adult ticks play a minor role in the transmission ecology of *B. burgdorferi* and are thus not explicitly modeled. Equations describe transitions between tick life stages and infection status from the modeling framework. (*b*.) The seasonal activity patterns of the tick developmental stages vary from near-continuous activity of all stages throughout the year (*i*) (Diuk-Wasser et al. 2006; Ogden et al. 2018) to developmental stages with temporally divergent activity seasons of short-duration within the US (iii) (Diuk-Wasser et al. 2006; Ogden et al. 2018). This latter tick-stage phenology (iii) is thought to result in high transmission of *B. burgdorferi* as large proportions of hosts are infected by nymphs (III) prior to larval activity (I). Black lines show larval activity and gray lines show nymhpal activity. Note that the larvae and nymphs that feed in the same summer are from different cohorts.

Here we develop a model to study the evolutionary ecology of parasite transmission given different phenological scenarios using the *B. burgdorferi-Ixodes* tick system as a natural example. The relative simplicity of our model makes mathematical analyses tractable while capturing the fundamental impact of phenology on parasite fitness. This impact unfolds over two time-scales: the within-season dynamics of infection, and the between-season demography of the vector (Bewick et al. 2016). Previous work (Dunn et al. 2013; Ogden et al. 2004) considered the within-season dynamics of infection but did not account for the between-season population dynamics of the vector species. The latter is an important factor as phenology can alter vector population sizes resulting in an ecological feedback impacting parasite fitness. Our analysis builds on a modeling framework that integrates these effects (Bewick et al. 2016) and demonstrates a general approach for studying both the short- and long-term impacts of vector phenology for parasite fitness. We use the Lyme disease system to describe our approach, although the modeling framework applies to all parasites that require multiple transmission events to complete their life cycle (*e.g*. West Nile Virus, Leishmania parasites, *Yersinia pestis*). Our framework can be extended to study how specific vector life history traits, such as differential mortality throughout the year, impact parasite fitness. Additionally, our straightforward framework makes further investigation of the evolutionary pressure imposed by phenology possible.

## Model

We model the transmission of *B. burgdorferi* between *I. scapularis* and a main vertebrate reservoir, the white-footed mouse, *Peromyscus leucopus* (LoGiudice et al. 2003). Our model tracks the within-season dynamics of nymphal and larval population activity and uses these dynamics to compute the between season changes in overall infection prevalence.

Within-season dynamics describe the duration of nymphal and larval emergence and feeding activity in continuous time from the beginning of each season (*t* = 0) to the end (*t* = *τ*). The life-cycle we model is depicted in Figure 1a. Ticks start their life-cycle uninfected, but may pick up the infection as larvae by feeding on an infected mouse (Magnarelli et al. 1987). Larvae then overwinter and emerge as nymphs in the next season who can transmit the infection to new mice who are also born uninfected (Hofmeister et al. 1999). The state variables *L*_•_(*t*), *N*_•_(*t*), and *M*_•_(*t*) represent larval, nymphal, and mouse populations, where the subscripts denote the host-seeking status of ticks (*q* for questing for a host or *f* for fed), as well as infection status of ticks and mice (*i* for infected, *u* for uninfected). Thus, *L_q_* denotes the questing larvae (who by definition cannot be infected), while *L_if_* denotes fed larvae that are infected. We make the common assumption that mortality and vital rates for both ticks and mice are not impacted by their infection status (Schwanz et al. 2011; Gage et al. 1995). The total mouse population size is *M* = *M_i_* + *M_u_*. The within season dynamics are given by the following system of ordinary differential equations:

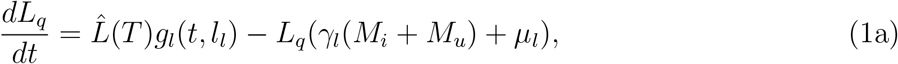

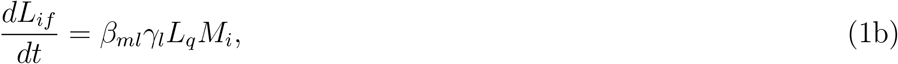

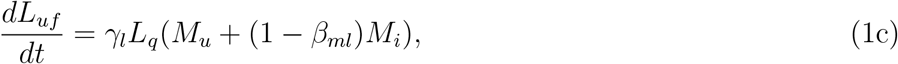

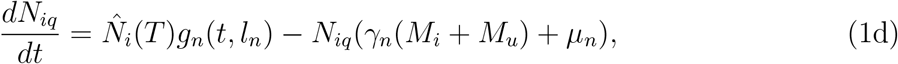

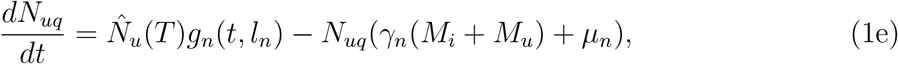

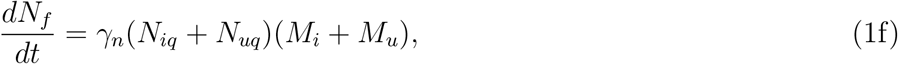

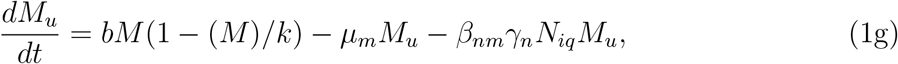

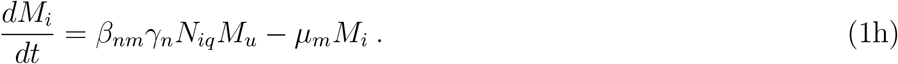

Here, 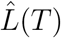 represents the total larval population to emerge in year *T*, as determined by the number of nymphs that have successfully fed in the previous year, *T* − 1, survived to adulthood, and reproduced (given by equation (5) below). Similarly, 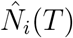 and 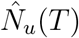 represent the total number of questing infected and uninfected nymphs that emerge in year *T* as determined by the number of infected and uninfected larvae at the end of the previous year and the probability of over-winter survival (see equations (3) and (4)). All other parameters are described in Table 1.

**Table 1:** Model parameters and their respective values. Time is measured in days for all parameters.

| Parameter | Description | Value |
| --- | --- | --- |
| $t_{\bullet 0}$ | start of activity period for tick life stage $\bullet$ | varies |
| $t_{\bullet f}$ | end of emergence period for tick life stage $\bullet$ | varies |
| $l_{\bullet}$ | length of emergence period for tick life stage $\bullet$ | varies |
| $\hat{L}$ | size of emerging larval population | varies |
| $\hat{N}_i$ | size of emerging infected nymphal population | varies |
| $\hat{N}_u$ | size of emerging uninfected nymphal population | varies |
| $\gamma_l$ | density dependent contact rate between larvae and mice | 0.004 (Randolph 1999a) |
| $\gamma_n$ | density dependent contact rate between nymphs and mice | 0.008 (Randolph 1999a) |
| $\beta_{nm}$ | transmission probability from nymphs to mice | 0.83 (Davis and Bent 2011) |
| $\beta_{hm}$ | transmission probability from mice to larvae | 0.6 (Davis and Bent 2011) |
| $\mu_l$ | larval death rate | 0.015 (Ogden et al. 2005a) |
| $\mu_n$ | nymphal death rate | 0.015 (Ogden et al. 2005a) |
| $\mu_m$ | mouse death rate | 0.01 (Schug et al. 1991) |
| $b$ | mouse birth rate | 0.1 (Schug et al. 1991) |
| $k$ | mouse carrying capacity | varies (Ostfeld et al. 1996b) |
| $\sigma_l$ | larval overwintering survival probability | 0.21 (Davis and Bent 2011) |
| $\sigma_n$ | compound fecundity and survival parameter | 10 (Davis and Bent 2011) |
| $\alpha$ | density dependence parameter | 0.0045 |
| $\tau$ | season length | 210 (Ogden et al. 2018) |

The functions *g_l_*(*t, l_l_*) and *g_l_*(*t, l_n_*) are probability density functions describing the timing and length of larval and nymphal emergence respectively. We describe tick emergence using a uniform distribution for analytical tractability.

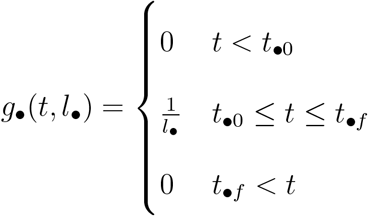

Where *t*_•0_ is the time emergence begins, *t*_•*f*_ is the time emergence stops and *l*_•_ is the length of the emergence period (*t*_•*f*_ – *t*_•0_ = *l*_•_). The uniform distribution establishes a constant emergence probability for ticks over *l*_•_ and thus spreads the emergence of the tick cohort evenly across the emergence period from *t*_•*f*_ < *t* < *t*_•0_. While our analysis relies on tick emergence following a uniform distribution, we conducted numerical simulations when tick emergence follows a Gamma distribution and found that the shape of the distribution does not qualitatively change our results (Appendix E.)

Equations (1a–1h) reduce to the following set of equations if we assume that the host population is at equilibrium, 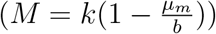:

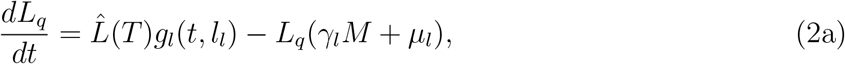

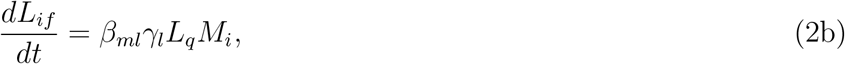

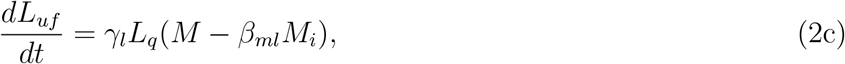

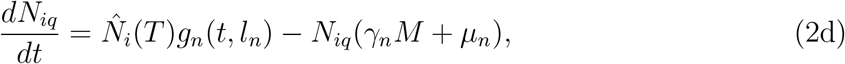

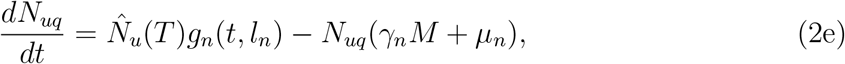

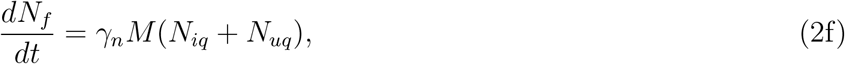

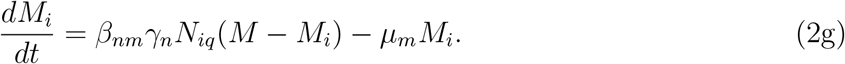

We solve equations (2a–2g) analytically in Appendix B.

### Between-season dynamics

The within-season dynamics described above are coupled to recurrence equations that describe the survival of larvae and nymphs between years. We do not follow the mouse population between years because the impact of overwintering infected mice on *B. burgdorferi* transmission is thought to be negligible (Bunikis et al. 2004; Anderson et al. 1987). The total number of infected and uninfected nymphs 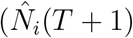 and 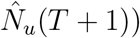 that emerge in a given year are given as a function of the number of infected and uninfected fed larvae at the end of the previous year (*L_if_*(*τ*) and *L_uf_*(*τ*)) as follows:

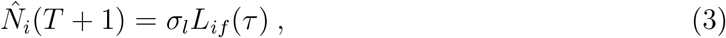

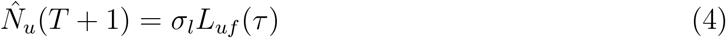

where *L_if_*(*τ*) and *L_uf_*(*τ*) are the infected and uninfected larval abundances at the end of the previous season (see Appendix B) and *σ_l_* is the larval overwintering survival probability.

Similarly, the total fed nymphal population at the end of the year *N_f_*(*τ*) gives rise to the population of larvae, 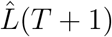, that emerges the following year as described by the map:

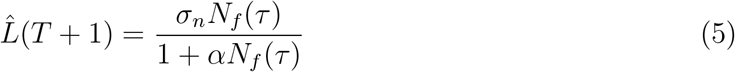

where *N_f_*(*τ*) is found by integrating (1e) over the season from (0, *τ*) as shown in Appendix B. *σ_n_* is the expected number of eggs produced per fed nymph, after accounting for survival to adulthood and for fecundity. The strength of density dependence on reproduction is determined by *α*.

With these functions, we can write the discrete, between season mapping of the total larval and nymphal abundances from one year to the next:

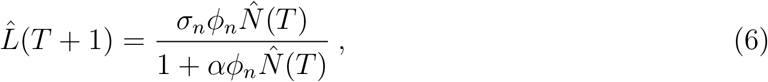

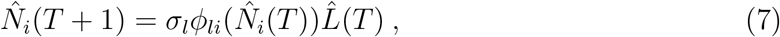

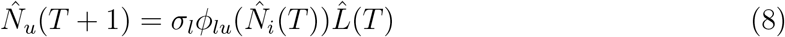

where *ϕ_n_* denotes the fraction of emerging nymphs that successfully feed as calculated from within-season dynamics (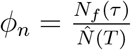; see Appendix A), and 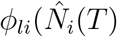) and 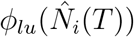 are functions of 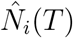 that denote the fraction of emerging larvae that become infected or remain uninfected through feeding as calculated from within-season dynamics (e.g., 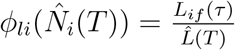; see Appendix B).

We next calculate the basic reproductive number, *R*_0_, to quantify the impact of phenology for *B. burgdorferi* fitness. *R*_0_ represents the average number of new infections caused by a single infected tick in an otherwise naïve population of mice and ticks (McCallum 2001), which gives the threshold for parasite invasibility given the phenology of both tick stages. *R*_0_ is computed as the number of infected nymphs that emerge in year *T* + 1 produced by a single infected nymph that emerged in year *T* in an otherwise uninfected population. Specifically, we consider a tick population that is at its demographic equilibrium without the infection, solved by setting 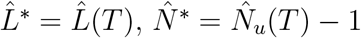, and 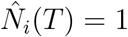 in equations (6)–(8). At this demographic equilibrium, *R*_0_ of a rare parasite infection is given as follows:

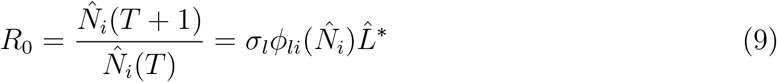

This *R*_0_ accounts for transmission between cohorts of ticks through intermediate mouse hosts in a given feeding season. When 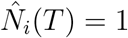, parasites persist in phenological scenarios where 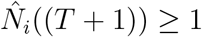 (*i.e*. slope is greater than or equal to unity). Details of the analytical approach are in Appendix C.

## Results

The rate of *B. burgdorferi* transmission from nymphs to mice to larvae is low in systems where either nymphs or larvae are continuously active (Fig. 2). Controlling for total population sizes, when nymphal feeding is evenly spread throughout the year, few nymphs feed at any given time, resulting in limited nymph-to-mouse transmission events. The proportion of infected mice remains constantly low as new infections occur at a similar rate as mouse mortality which replaces older, potentially infected mice with uninfected juveniles. Larval ticks rarely encounter infected mice, thus limiting mouse-to-tick transmission events. By contrast, seasonal nymphal activity concentrates nymph-to-mouse transmission events in time, causing a seasonal peak in mouse infection prevalence that decays due to mouse population turnover (Fig. 2). The duration of the nymphal activity period is negatively correlated with the rate at which infected mice accumulate as well as the maximum mouse infection prevalence (*e.g*. small *l_n_* values in Fig. 2). That is, nymphal activity periods of greater duration result in a lower maximum mouse infection prevalence that peaks later in the season (Fig. 2). Larval ticks that feed at or around the peak in mouse infection prevalence are more likely to encounter an infected mouse and to acquire *B. burgdorferi* before molting to nymphs.

**Figure 2:**
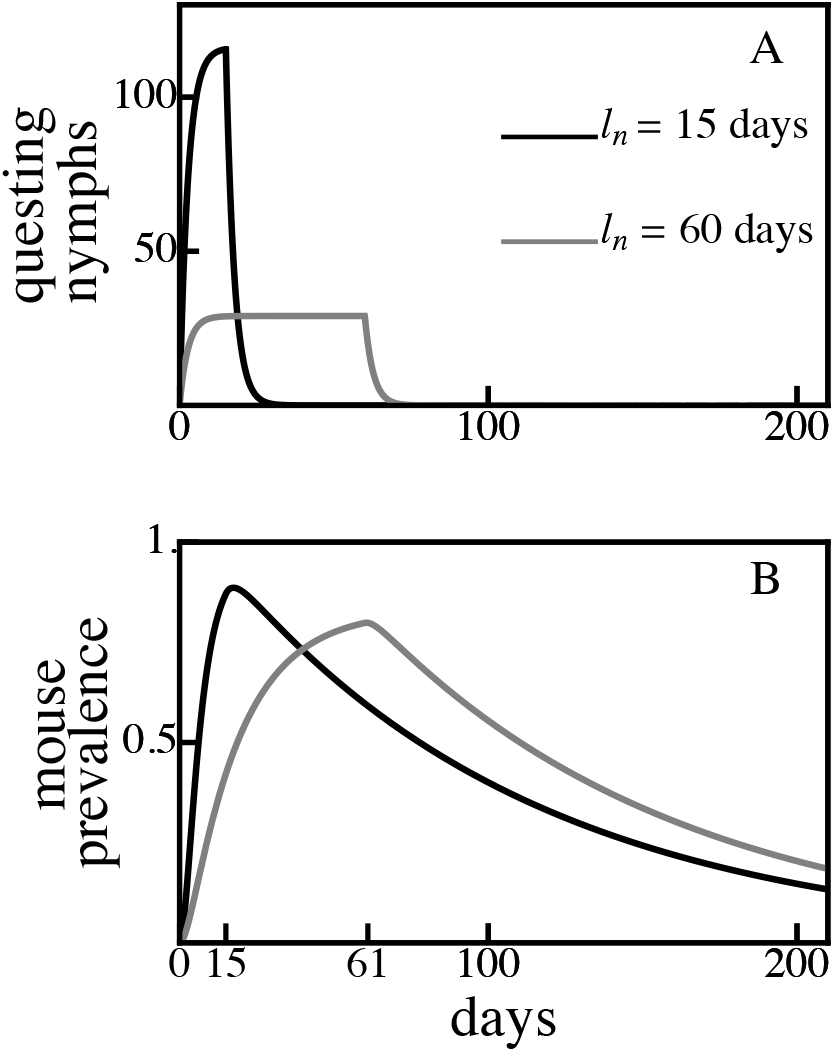
Concentrated nymphal emergence durations (*l_n_* = 15 days, *A*) result in a higher and earlier mouse infection prevalence peak (*B*) compared to longer nymphal emergence durations (*l_n_* = 60 days) for the same total tick population sizes. For example, a nymphal activity duration of 15 days (*l_n_* = 15 days, *A*) results in peak mouse infection prevalence (*B*) occurring on day 15 while *l_n_* = 60 days results in peak mouse infection prevalence occurring on day 61. In both models, 25% of emerging nymphs are infected, *k* = 50, *M* = *k*(1 – *μ_m_*/*b*) = 45. All other parameter values are shown in Table 1.

The fitness of *B. burgdorferi*, quantified by the basic reproductive number (*R*_0_), is greatest when larval activity is concentrated around the peak in the mouse infection prevalence, thus increasing the probability that each larva will feed on an infected mouse (Fig. 3A. and Fig. 4A). Larvae that are active long after the end of the nymphal activity period are likely to feed on an uninfected mouse due to decays in mouse infection prevalence caused by mouse population turnover (Fig. 3B and Fig. 4B). Similarly, larval activity periods that begin prior to nymphal activity periods result in the majority of larvae feeding on uninfected mice that have not acquired an infection from a feeding nymph.

**Figure 3:**
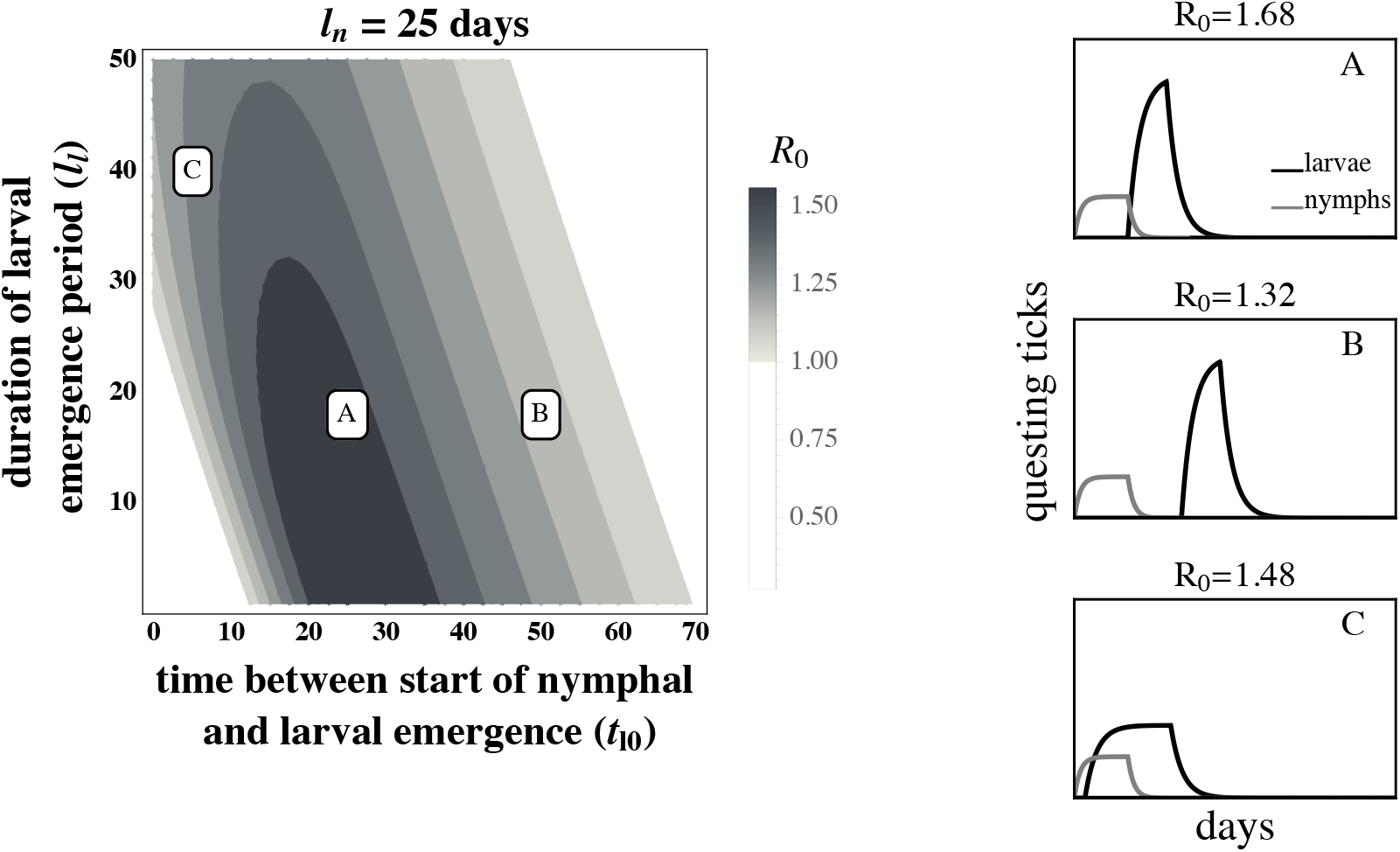
The basic reproductive number, *R*_0_, of *B. burgdorferi* is greatest when larval activity is concentrated around peak mouse infection prevalence. The left panel depicts *R*_0_ as a function of the duration of larval emergence (*l_l_*) and time between nymphal and larval emergence (*t*_*l*0_). Panels on the right depict within-season dynamics for representative timing parameter values indicated by their respective letters on the left panel. (*A*) Concentrated larval emergence (small *l_l_*) that begins slightly after nymphal emergence (20 < *t*_*l*0_ < 35) increases the probability that questing larvae feed on mice recently infected by nymphs (*t*_*l*0_ = 25, *l_l_* = 18). (*B*) Transmission decreases as larvae emerge later (*t*_*l*0_ > 35) because the larval cohort feeds after peak mouse infection prevalence (*t*_*l*0_ = 50, *l_l_* = 18). (*C*) When larval and nymphal emergence is more synchronous (small *t*_*l*0_), transmission to larvae increases as larval emergence duration increases (large *l_l_*) because more larvae feed after infectious nymphs (*t*_*l*0_ = 5, *l_l_* = 40). *B. burgdorferi* is not maintained in systems where *R*_0_ < 1. *R*_0_ is calculated assuming tick emergence is uniformly distributed (*U*(*l_l_*) where *l_l_* is the larval emergence duration, see Appendix C). 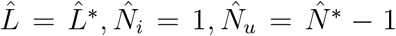 (see Appendix A.) *l_n_* = 25 days; all other parameter values are shown in Table 1.

**Figure 4:**
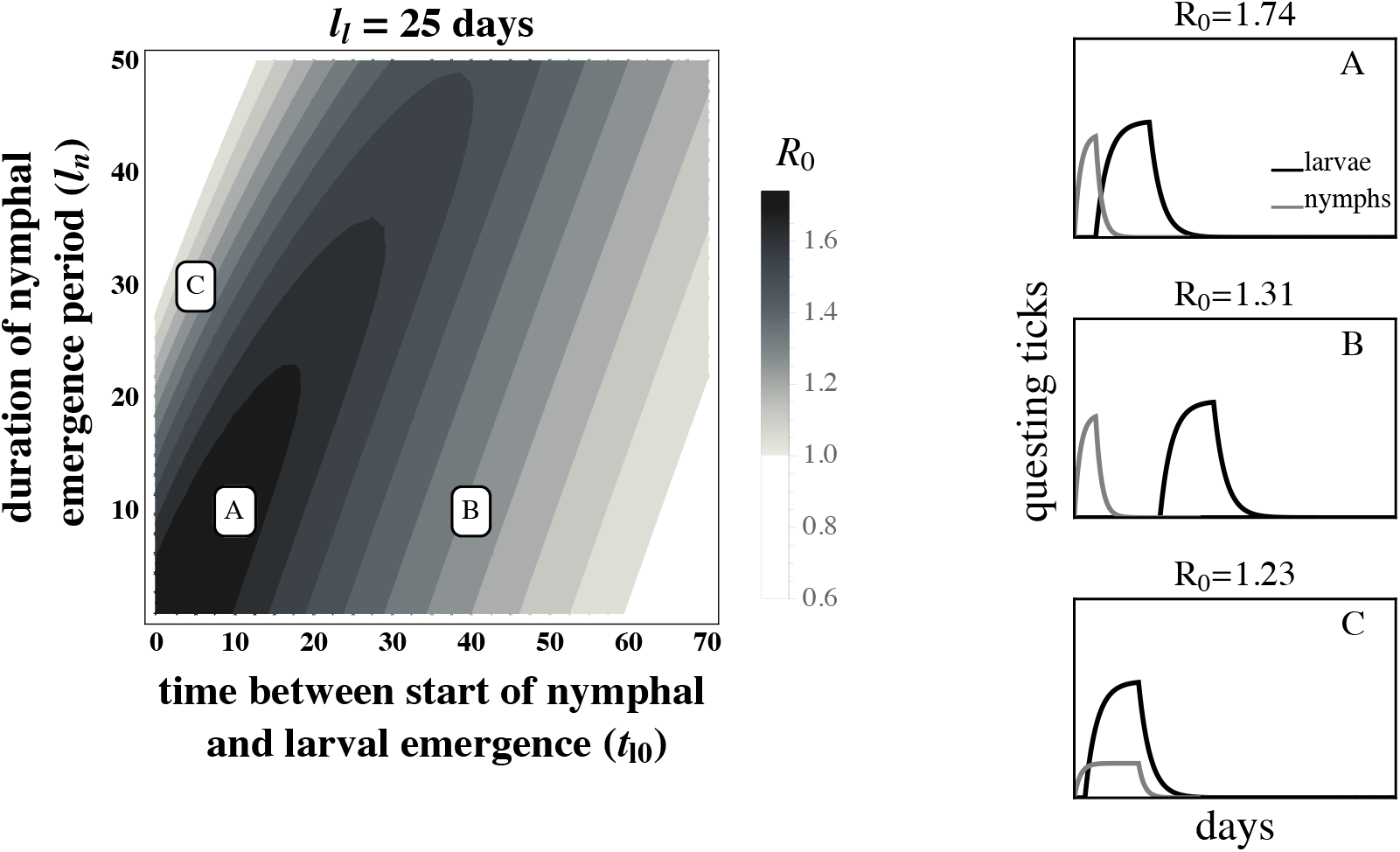
The basic reproductive number, *R*_0_, of *B. burgdorferi* is greatest when larval emergence begins shortly after nymphal emergence such that larvae feed during peak mouse infection prevalence. The left panel depicts *R*_0_ as a function of the time between the start of nymphal and larval emergence (*t*_*l*0_) and the duration of the nymphal emergence period (*l_n_*). The letters within the left panel indicate the parameter values used to depict representative within-season dynamics in the right panels. (*A*) Concentrated nymphal emergence (small *l_n_*) coupled with slight differences in nymphal and larval emergence time (*t*_*l*0_ < 10) increases the probability that questing larvae feed on mice infected by nymphs (*t*_*l*0_ = 10, *l_n_* = 10). (*B*) Longer durations between nymphal and larval emergence time (*t*_*l*0_ > 10) results in lower mouse-to-larvae transmission rates as many mice infected by nymphs die and are replaced by mice born uninfected such that larvae are likely to feed on uninfected mice (*t*_*l*0_ = 40, *l_n_* = 10). (*C*) Synchronous emergence (*t*_*l*0_ = 0) can also reduce *B. burgdorferi* fitness when nymphal emergence duration is long (large *l_n_*) as many larvae feed before mice become infected by nymphs (*t*_*l*0_ = 5, *l_n_* = 30). *R*_0_ is calculated assuming tick emergence is uniformly distributed (*U*(*l_n_*) where *l_n_* is nymphal emergence length, see Appendix C). *l_l_* = 25, 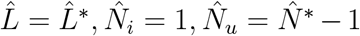 (see Appendix A). All other parameter values are shown in Table 1.

The effect of larval emergence duration depends on whether or not larval emergence coincides with nymphal emergence: concentrated larval emergence decreases *R*_0_ when larval and nymphal emergence periods are synchronous (Fig. 5) because most larvae feed before nymphs have a chance to infect the mouse population. Conversely, concentrated larval emergence tends to increase *R*_0_ when larvae emerge later than nymphs (Fig. 6). This occurs because nymphal emergence that slightly precedes larval emergence results in high mouse infection prevalence when larvae begin emerging (Fig. 6A), and concentrated emergence results in most larvae feeding when the prevalence of infection is still high. In both cases, *R*_0_ decreases with very broad larval emergence due to mouse turnover (Fig. 3C, Fig. 4C, Fig. 6B, Fig. 5B).

**Figure 5:**
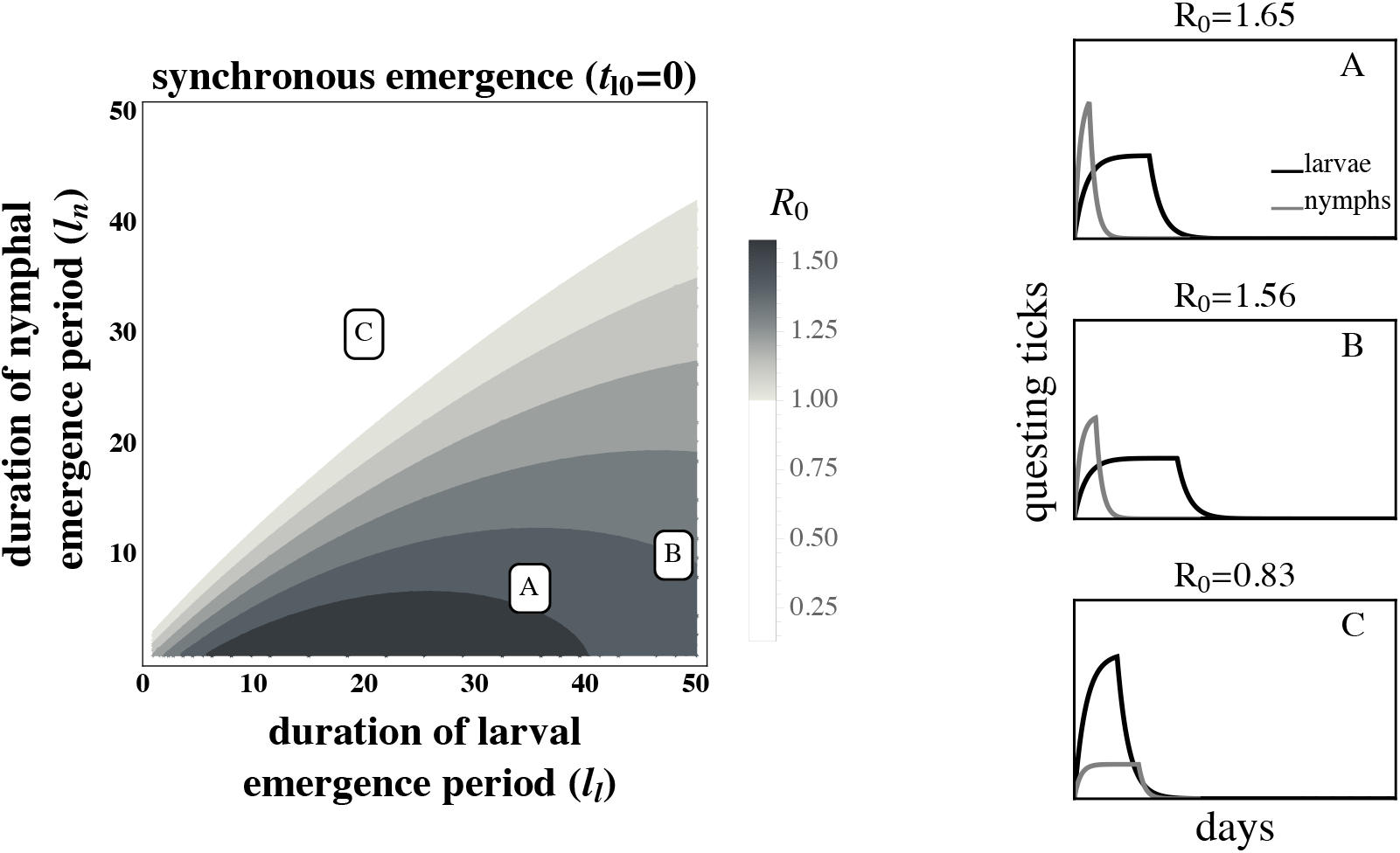
The larval emergence duration that maximizes *R*_0_ for *B. burgdorferi* is conditioned on nymphal emergence duration. *R*_0_ is high if larval emergence duration is slightly longer than nymphal emergence duration (*l_l_* > *l_n_* in (*A*) and (*B*)), thus allowing larvae to feed on mice that were previously fed upon by nymphs. However, *R*_0_ decreases when larval emergence duration is much longer than nymphal emergence duration (*R*_0_ of (*B*) < *R*_0_ of (*A*)) as late emerging larvae can feed on uninfected mice born after the nymphal activity period. Transmission from mice to larvae is low when the larval emergence duration is less than the nymphal emergence duration (*l_l_* < *l_n_* in (*C*)) because many larvae feed before infectious nymphs. *B. burgdorferi* is not maintained in systems where *R*_0_ < 1. *R*_0_ is calculated assuming tick emergence is uniformly distributed (*U*(*l_l_*) where *l_l_* is the larval emergence length. See Appendix C for details). 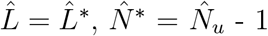 (see Appendix A). *t*_*l*0_ = 0, (*A*) *l_l_* = 35, *l_n_* = 7 (*B*) *l_l_* = 48, *l_n_* = 10 (*C*) *l_l_* = 20, *l_n_* = 30. *t*_*l*0_ = 0 days; all other parameter values are shown in Table 1.

**Figure 6:**
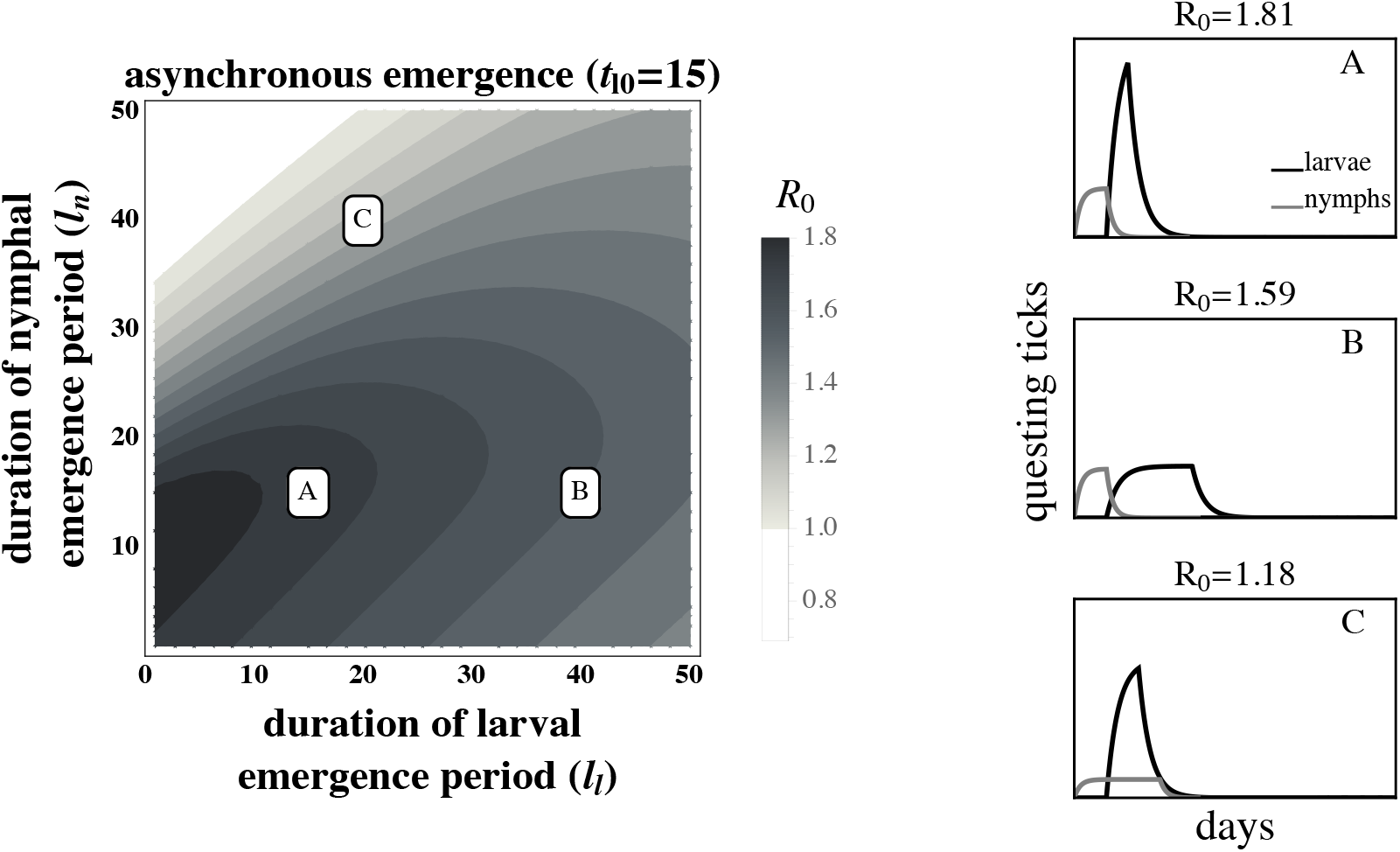
Highly concentrated larval emergence increases *R*_0_ when larvae emerge slightly after nymphs. (*A*) Concentrated nymphal emergence drives high mouse infection prevalence and results in high transmission to larvae when larval emergence is tightly concentrated (*l_l_* = 10, *l_n_* = 15). (*B*) Transmission from mice to larvae decreases as larval emergence duration increases because larvae are more likely to feed on uninfected mice born after nymphal activity (*l_l_* = 40, *l_n_* = 15). (*C*) Transmission from mice to larvae also decreases if larval emergence duration is highly concentrated and nymphal emergence duration is broad because many larvae feed before nymphs infect mice (*l_l_* = 15, *l_n_* = 40). *R*_0_ is calculated assuming tick emergence is *U*(*l_l_*) where *l_l_* is the larval emergence length (see Appendix C.) 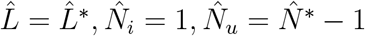 (see Appendix A.) *t*_*l*0_ = 15 days; all other parameter values are shown in Table 1.

## Discussion

Phenology is a fundamental component of all ecological interactions. Interactions between organisms such as competition, predation, and parasitism are predicated on temporal overlap of interacting species or life-stages. Similarly, host or vector phenology impacts parasite fitness by temporally structuring transmission events between interacting hosts or life stages. Host or vector phenological patterns can even determine whether a pathogen is highly-abundant or is unable to persist (Fig. 3). The ubiquity of seasonal activity among hosts and vectors, as well as the geographic variation in seasonal activity patterns, underscores the importance of phenology for the distribution and abundance of many pathogenic microbes including malaria, rabies, tapeworm, and Lyme disease (Hoshen and Morse 2004; Gremillion-Smith and Woolf 1988; Anderson 1974; Ogden et al. 2018). Here we derive the basic reproductive number, *R*_0_, for a transmission dynamics model that explicitly considers the impact of phenology on both parasite transmission and vector demography to assess the impact of vector phenology on parasite transmission and fitness using the Lyme disease system as an example. Our results are benchmarked by field data that show a link between the regional differences in tick phenology and differences in *B. burgdorferi* distribution and abundance (Ogden et al. 2018). Investigation of parameter space in this model revealed the novel insight that large temporal differences between the activity periods of tick-life stages decreases *B. burgdorferi* fitness.

Our model accounts for an important ecological feedback between vector demography and parasite fitness by incorporating the impact of phenology on demography. This is achieved by capturing both within-season infection dynamics and between-season vector demography in our mathematical analysis. Tick population sizes increase with earlier and more concentrated emergence because most ticks have sufficient time to successfully find a host before the season ends. By contrast, late or less concentrated emergence results in fewer ticks feeding before the season ends as the time available for later-emerging ticks to find a host is effectively shortened. The impact of this demographic feedback is limited at high mouse density but increases rapidly at low mouse density (see Appendix D). Extensions to this framework where vector mortality or contact rates with hosts vary throughout the year are likely to exacerbate the impact of phenology on demography. These results suggest that disregarding between-season demographic dynamics could under-estimate parasite fitness (*R*_0_) when ticks emerge early and over-estimate *R*_0_ when ticks emerge later. The importance of this ecological feedback is reflected in the finding from a next-generation model focusing on within-season (but not between-season) impacts of tick phenology on *B. burgdorferi* fitness which indicated that vector demography is one of the most important model parameters affecting *R*_0_ (Dunn et al. 2013).

Parasite fitness is maximized when the activity periods of vector life stages are of short duration (Fig. 2). Continuous nymphal activity temporally distributes the finite number of nymph-to-mouse transmission events such that mice become infected at a low rate throughout the season. Mouse infection prevalence remains continually low because mice that die, including infected mice, are replaced by uninfected juveniles at rates similar to the rate at which new infections are introduced. Mouse-to-larvae transmission events are similarly rare as most larvae feed on the relatively abundant uninfected mice. By contrast, seasonal nymphal activity concentrates nymph-to-mouse transmission events leading to many new mouse infections over a short period of time. Mouse infection prevalence increases rapidly during the nymphal activity period, as new infections occur at a much greater rate than mouse mortality, and subsequently decline when new infections stop at the end of the nymphal activity period (Fig. 2). Transmission from mice to larvae is very high if larval activity coincides with high mouse infection prevalence (Fig. 3A. and Fig. 4A.) The temporal concentration of infected hosts is likely to have important consequences for the transmission success and fitness of most pathogens (Altizer et al. 2006; Martinez 2018).

Extended periods between nymphal and larval activity results in limited transmission efficiency (Fig. 3B and Fig. 4B). This novel prediction for the Lyme disease system is caused by the decay in mouse infection prevalence following nymphal activity due to mouse mortality and the birth of uninfected mice (Hofmeister et al. 1999; Wright et al. 1990). Thus, larvae feeding long after the nymphal activity period have a greater probability of feeding on uninfected mice than those that feed shortly after the nymphal activity period. While high mouse turnover is the norm in this system (Schug et al. 1991), lower mouse turnover would extend the period of high mouse infection prevalence and moderate the declines in parasite fitness caused by extended periods between larval and nymphal emergence.

Parasite fitness is predicted to be greatest when all individuals within each developmental stage feed simultaneously and larvae feed immediately after nymphs. This result relies on the assumption that there is no limit to the number of ticks that can feed on a mouse at any given time. Realistically, the number of ticks per mouse is limited by grooming and foraging behaviors. Incorporating a maximum number of ticks per mouse will alter the prediction that simultaneous emergence within life stages maximizes parasite fitness as most ticks will fail to find an available host, resulting in fewer fed ticks each year and thus a lower *R*_0_. Further, accounting for spatial aggregation of host-seeking larvae would increase the impact of a maximum number of ticks per mouse (Ostfeld et al. 1996c,a, 2018; Devevey and Brisson 2012). Incorporating this ecological realism will cause intermediate emergence concentrations to result in more infected larvae.

The observed fitness of *B. burgdorferi* in different Lyme disease foci in North America corresponds qualitatively with model predictions. For example, the relatively continuous activity of both tick developmental stages in the Southeastern United States has been proposed as a factor leading to the relatively low *B. burgdorferi* fitness observed in the region. In the Midwestern United States, where larvae and nymphs are synchronously active during a limited period, *B. burgdorferi* transmission is lower than in the Northeastern US but much greater than where both stages are more continuously active (Fig. 5) (Hamer et al. 2012). The correlation between *B. burgdorferi* fitness observed in nature and the expected fitness differences given the observed phenological patterns suggests that both the duration of seasonal activity and the relative timing of activity periods may impact transmission success and parasite fitness (Figs. 3, 4, 6, 5). However, vector phenology is unlikely the only cause of the differences in *B. burgdorferi* transmission success among these regions as many other features that are known to impact *B. burgdorferi* also differ including host community composition, tick host preferences, and landscape and climatic features (James and Oliver Jr 1990; LoGiudice et al. 2003; Brisson and Dykhuizen 2004; Ogden et al. 2005b; Brisson et al. 2008; Khatchikian et al. 2012; Vuong et al. 2014; Adalsteinsson et al. 2016; Vuong et al. 2017; Adalsteinsson et al. 2018). Nevertheless, our results add to the body of literature that suggests tick phenology can impact *B. burgdorferi* fitness.

Our model captures the impact of phenology on *B. burgdorferi* transmission and fitness in a much simpler modeling framework than previously published studies that successfully address several hypotheses specific to the system (Ogden et al. 2004, 2008; Dunn et al. 2013). In particular, previous work focused on accurately predicting *B. burgdorferi* incidence given phenological scenarios in several realistic environments that depend upon several dozens of parameters, all of which require empirical validation (Ogden et al. 2004). In contrast, our model has 15 parameters and a straightforward structure. This relative simplicity allows our model to serve as a basis for studying phenological impacts in a broad range of environmental scenarios and disease systems as well as exploring the ramifications of other complicating factors such as the evolutionary dynamics of virulence.

As all disease systems exhibit seasonality, phenological drivers may have large impacts on the transmission success, and disease risk from, many parasites. Geographic variation in host or vector phenology may also be an important driver of documented variations in pathogen prevalence and disease risk (Altizer et al. 2006; Martinez 2018). Public health predictions of disease risk may be improved by accounting for phenological variation. Further, the dramatic shifts in host and vector phenology driven by global climate change (Penuelas 2001; Meyer et al. 2014; Post et al. 2001; Johansson et al. 2014) may result in equally dramatic shifts in pathogen prevalence at regional or global scales.

## Declarations

### Funding

This work was supported by the National Institutes of Health (T32AI141393 (HM) R01AI142572 (DB), R01AI097137 (DB)); the National Science Foundation (DEB-1354184 (DB)); and the Burroughs Wellcome Fund (1012376 (DB)).

### Conflicts of interest/Competing interests

The authors declare that they have no conflict of interest.

### Ethics approval

Not applicable

### Consent to participate

Not applicable

### Consent for publication

All authors gave final approval for publication.

### Availability of data and material

Not applicable

### Code availability

Mathematica code written to generate figures in the main text and appendix are included as Supporting Information.

### Author’s contributions

Hannelore MacDonald and Dustin Brisson conceived of the presented idea; Hannelore MacDonald, Erol Akçay, and Dustin Brisson developed the theoretical framework; Hannelore MacDonald conducted the mathematical analysis and performed numerical simulations; Erol Akçcay supervised the mathematical analysis and numerical simulations; All authors wrote the manuscript and gave final approval for publication and agree to be held accountable for the work performed therein.

## Author’s contributions

HM and DB conceived of the presented idea; HM, EA, and DB developed the theoretical framework; HM conducted the mathematical analysis and performed numerical simulations; EA supervised the mathematical analysis and numerical simulations; HM, EA and DB wrote the manuscript; All authors gave final approval for publication and agree to be held accountable for the work performed therein.

This work was supported by the National Institute of Health (T32AI141393 (HM) R01AI142572 (DB), R01AI097137 (DB)); the National Science Foundation (DEB-1354184 (DB)); and the Burroughs Wellcome Fund (1012376 (DB)).

## Appendix A

In Appendix A we derive between-season equilibrial solutions for tick demography (A.1a – A.1d), ignoring infection status. The following differential equations describe within-season tick population dynamics, valid from (0, *τ*) where *τ* is the length of the tick feeding season. For simplicity, we assume that the mouse population is constant so that 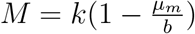 where *b* is the mouse birth rate, *k* is the carrying capacity and *μ_m_* is the mouse death rate.

All other parameters are the same as described in the main text.

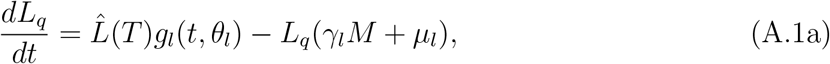

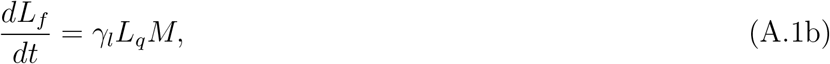

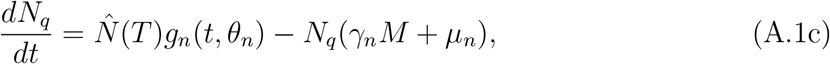

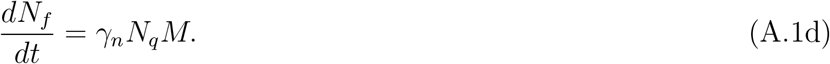

(A.1a–d) is solved analytically by describing tick emergence using a uniform distribution

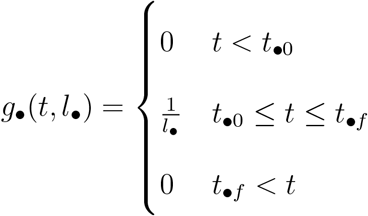

Within-season dynamics are coupled to recurrence equations that describe nymphal and larval survival between years. The total fed larval population at the end of the year, *L_f_*(*τ*) gives rise to the population of nymphs 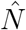 that will emerge the following year, described by the map

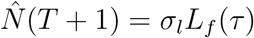

Where *σ_l_* accounts for the survival between fed larvae and questing nymphs and the number of fed larvae at time *τ* is given by

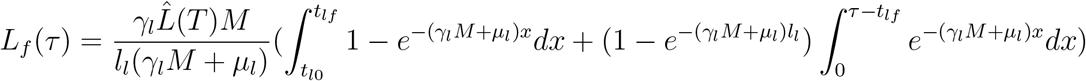

Similarly, the total fed nymphal population at the end of the year, *N_f_*(*τ*), gives rise to the population of larvae 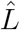 that will emerge the following year, described by the map

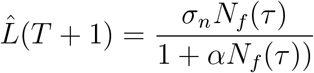

This expression accounts for the expected number of questing larvae produced per nymph that feeds to repletion after accounting for survival through adulthood, density dependent adult fecundity and survival from egg to questing larva. The number of fed nymphs at time *τ* is given by

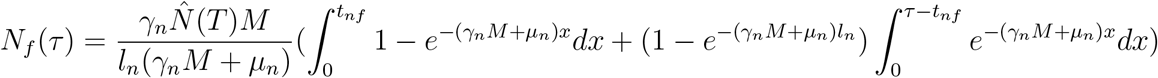

If we define

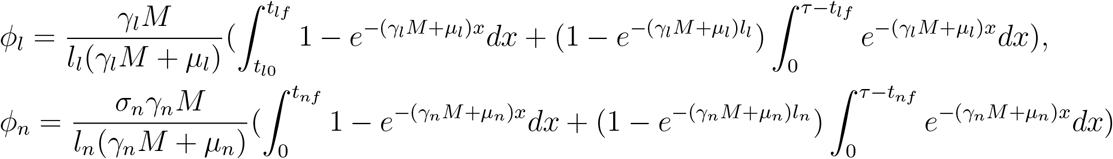

The maps for 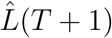 and 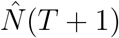 can be written as

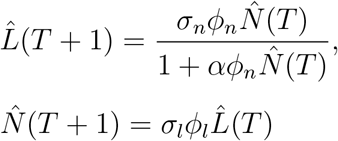

The equilibrium population size 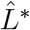 is then

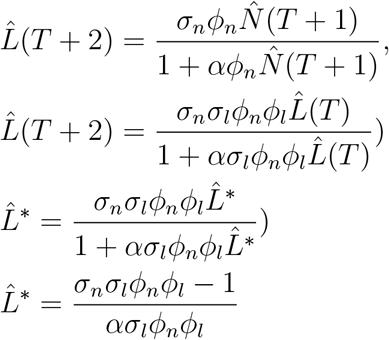

Similarly for 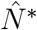

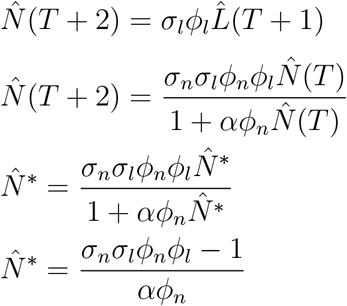

The stability of these equilibrium points is found by considering the biennial maps of the tick life cycle

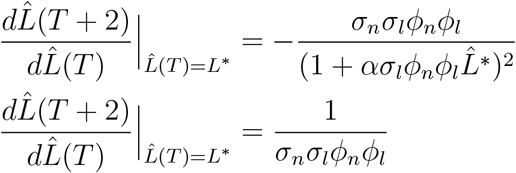

This system is stable for *σ_n_σ_l_ϕ_n_ϕ_l_* > 1. *σ_n_, ϕ_n_*, and *ϕ_l_* are always less than 1. Therefore, the stability of the tick demographic equilibrium depends on maintaining 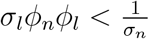. Figures in the main text assume both tick life-stages are at equilibrium by using *L*\* and *N*\* for the sizes of emerging tick cohorts at the beginning of the season. 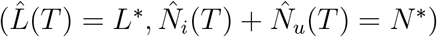.

## Appendix B

In Appendix B we derive analytical solutions for Equations (2a–2g) from the main text. Equations (2a–2g) assume that the host population is at equilibrium, 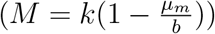. We put these equations again in Appendix B:

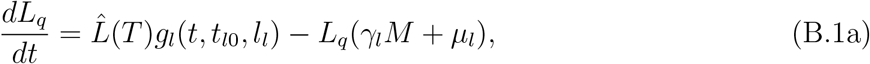

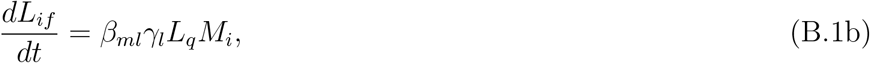

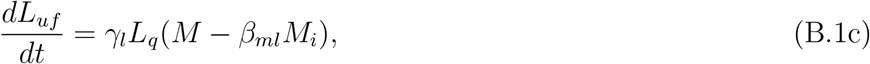

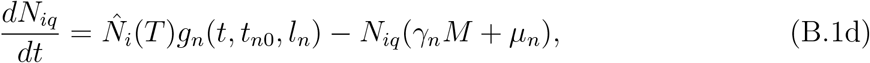

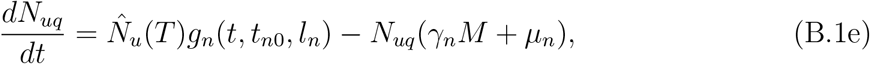

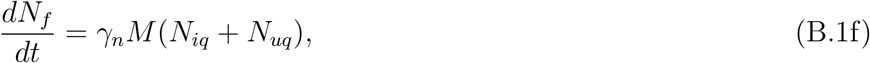

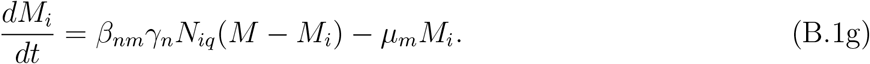

The system (B.1a–g) is solved analytically by describing tick emergence using a uniform distribution

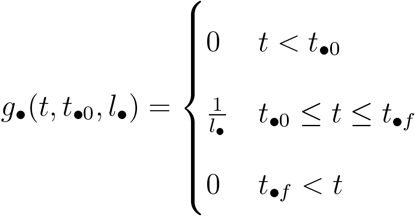

where *t*_•0_ denotes the start of emergence, *l*_•_ denotes the length of emergence and *t*_•*f*_ denotes the end of emergence (*t*_•0_ + *l*_•_ = *t*_•*f*_). The season begins with the emergence of the nymphs (*t*_*n*0_ = 0). Larval emergence, *t*_*l*0_ can begin concurrently with nymphal emergence (*t*_*l*0_ = 0) or have a start time that is offset relative to nymphs (*t*_*l*0_ > 0).

To solve for the within-season dynamics we first find the time-dependent solutions for questing and fed nymphs. Emerging nymphs are split by their infection status 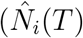 and 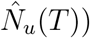 which was determined by whether they were infected during their bloodmeal as larvae in the previous season.

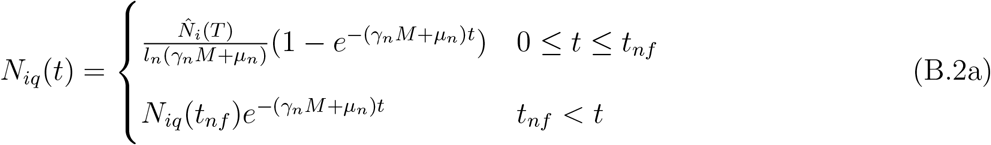

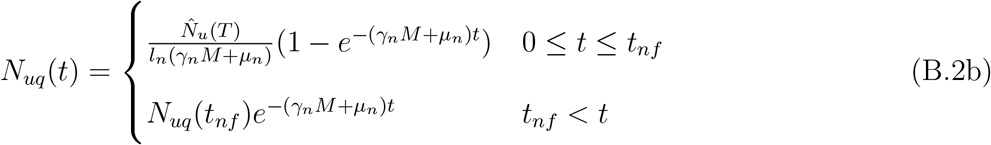

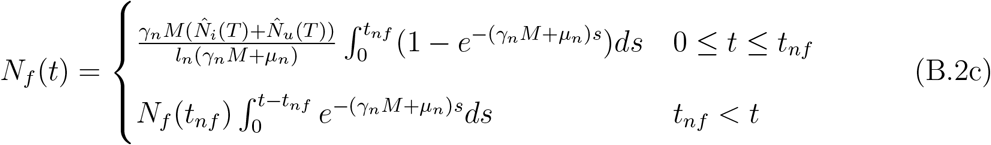

We then use *N_iq_*(*t*) to find the solution for mouse infection dynamics:

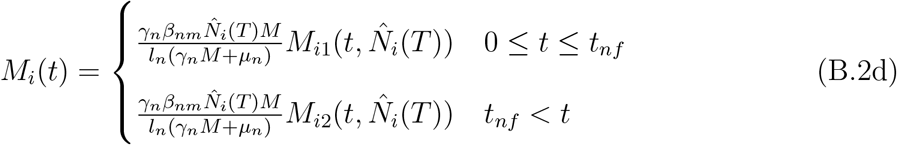

(B.2d) depends on the activity of questing nymphs and is split by whether nymphs are emerging, *M*_*i*1_(*t*) for 0 ≤ *t* ≤ *t_nf_* or have finished emerging, *M*_*i*2_(*t*) for *t_nf_* < *t* < *τ*.

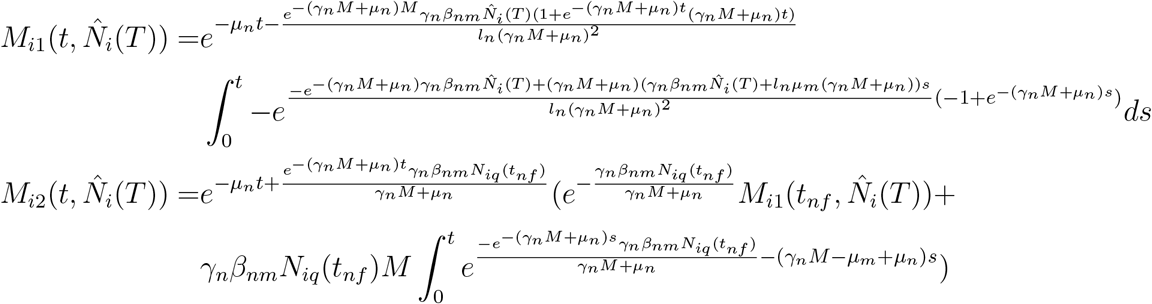

We next find the time-dependent solutions for questing and fed larvae, where 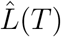 denotes the emerging larval cohort. We use the solution for *M_i_*(*t*) to split fed larvae by whether they became infected while feeding on a mouse.

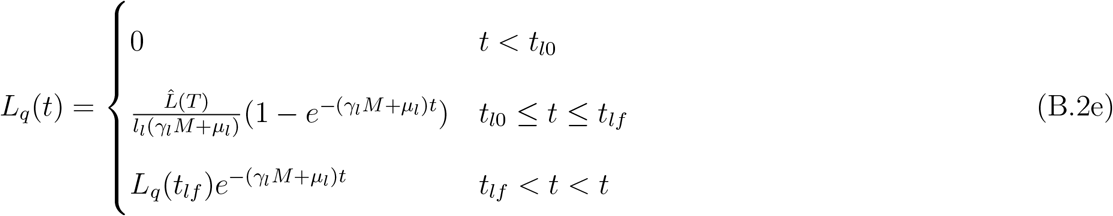

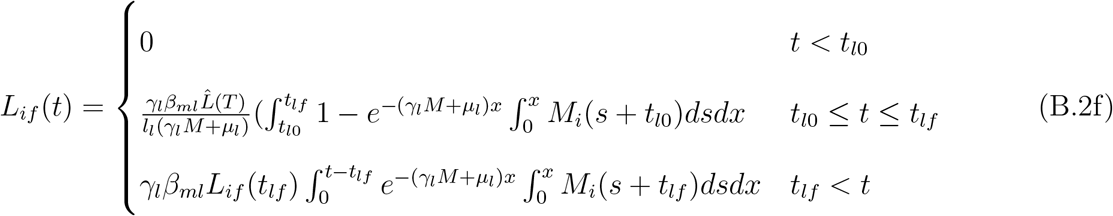

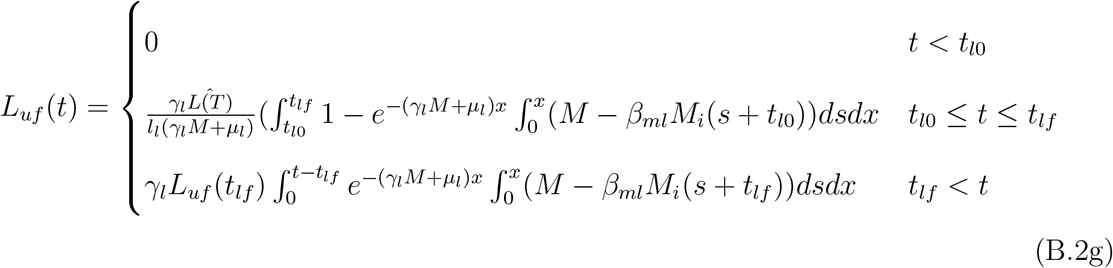

The total number of fed infected larvae by the end of the season, *L_if_*(*τ*), fed uninfected larvae by the end of the season, *L_uf_*(*τ*) and total fed larvae by the end of the season *L_f_*(*τ*) are given by

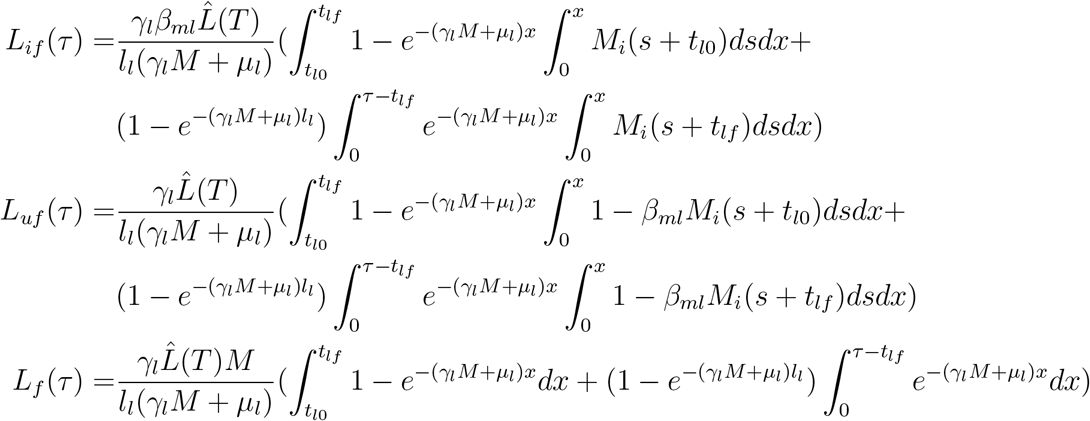

Note that both *L_if_*(*τ*) and *L_uf_*(*τ*) are dependent on 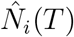 through the transmission dynamics of *M_i_*(*t*).

The total number of fed nymphs by the end of the season is given by

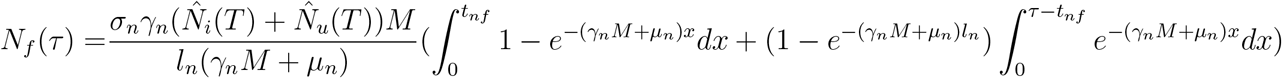

We can also write *L_if_*(*τ*), *L_uf_*(*τ*) and *N_f_*(*τ*) in terms of the total number of emerging ticks for a given season, 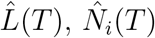, and 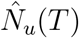.

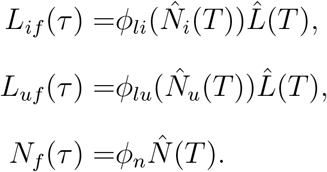

Where *ϕ_n_* denotes the fraction of emerging nymphs that feed over a season as calculated from within-season dynamics and 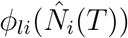 and 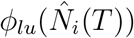 are functions of 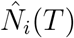 that denote the fraction of emerging larvae that feed and become infected or remain uninfected respectively, as calculated from within-season dynamics.

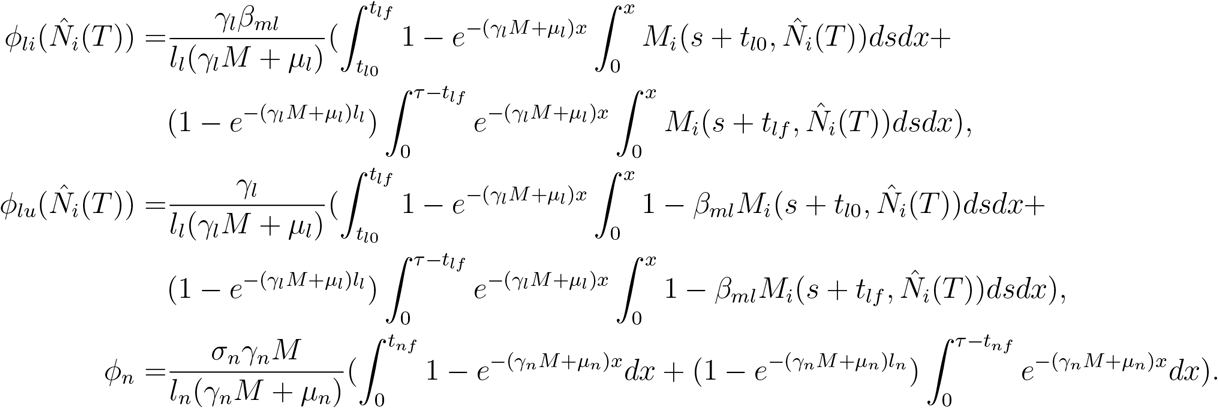

Discrete annual maps of each population can then be written as

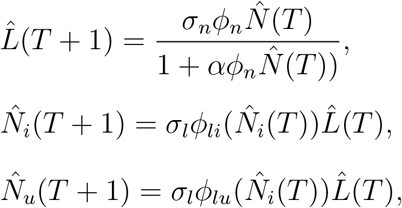

To check the stability of tick populations, consider the biennial maps

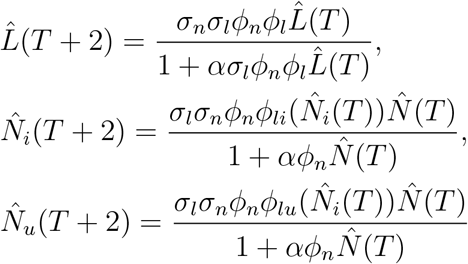

Infection status does not impact demographic rates. Therefore the larval equilibrium size 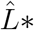 and total nymphal equilibrium size 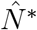 in the infection subsystem are identical to the result found above in Appendix A that ignores infection. 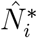 and 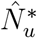 are both stable for the same conditions given in Appendix A because they are upper bounded by 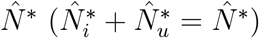. Assuming the stability conditions in Appendix A are met, the nontrivial solutions for 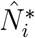 and 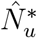 are unique because *ϕ_li_, ϕ_lu_* < 1 and the transmission terms are 0 < *β_nm_, β_ml_* < 1.

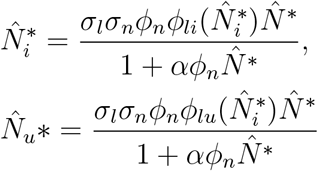

## Appendix C

In Appendix C we derive *R*_0_ to study how phenology impacts thresholds for parasite persistence. *R*_0_ is computed as the number of infected nymphs that emerge in year *T* + 1 produced by a single infected nymph that emerged in year *T* in an otherwise uninfected population. Specifically, we consider the stabiliy of the disease-free equilibrium when a rare infected nymph is introduced into the tick population, solved by setting 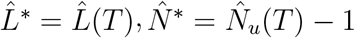, and 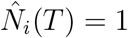 in equations (6) and (8).

Three distinct cases of phenological patterns are relevant to this system: (1) Emergence of both tick stages overlap and larvae finish emerging before nymphs finish emerging (2) Emergence of both tick stages overlap and nymphs finish emerging before larvae finish emerging (3) Nymph emergence ends before larvae emergence begins. Each case needs to be analyzed separately to account for the time dependent differences in the dynamics.

Using Case 1 as an example to sketch our derivation:

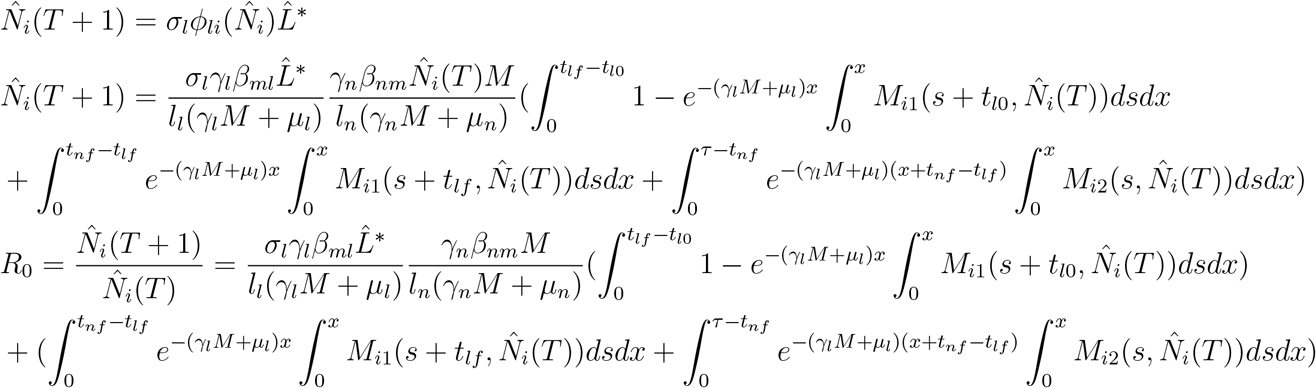

for 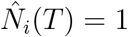, parasites persist in phenological scenarios where 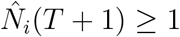. Parasite fitness is maximized when peak larval activity coincides with peak host prevalence. When *R*_0_ > 1, the number of infected nymphs, 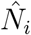 reaches a stable T-periodic equilibrium

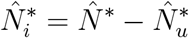

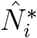 for a given phenological scenario can be found by solving for the value of 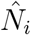 that satisfies *R*_0_ = 1. Again, using Case 1 as an example:

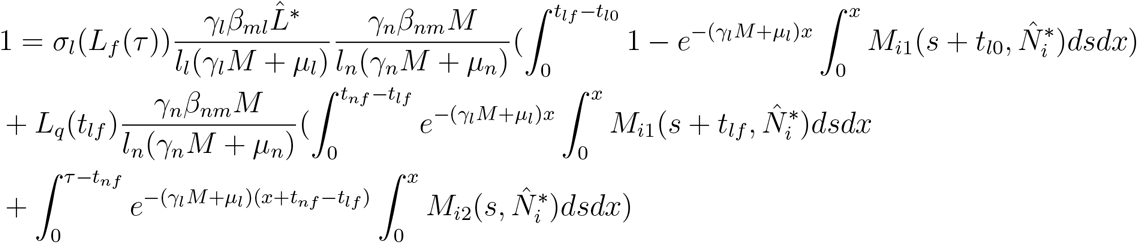

If the total tick population is stable (see Appendix A), 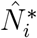 is upper bounded by 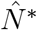 and is therefore stable as well.

## Appendix D

In Appendix D we demonstrate that tick phenological patterns can alter tick equilibrium population sizes and impact *R*_0_. Not accounting for the impact of phenology on between-season tick population demography by assuming constant tick populations each year leads to over or under estimates of tick demography (Figure A-1) and parasite fitness (Figures A-2, A-3). Our model in contrast accounts for the feedback between vector demography and parasite fitness by considering both within-season transmission and vector population dynamics (see Appendix B and C) and between-season vector demography (see Appendix A and B). Mouse density impacts *R*_0_ (Figure A-4); the error in *R*_0_ estimate when not accounting for population dynamics is especially high when mouse density is low.

*R*_0_ is a function of the equilibrium larval population size, 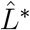, which is in turn determined by tick phenology (see equation (A.3) in Appendix A).

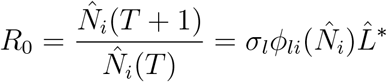

The ecological feedback between tick phenology, tick demography and *R*_0_ is ignored if a constant 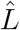 is assumed for all phenological patterns.

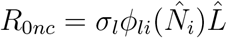

where *R*_0*nc*_ refers to the fact that *R*_0_ is “not corrected.”

The relative error between *R*_0_ and *R*_0*nc*_ is calculated as 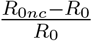.

## Appendix E

In Appendix E we present numerical simulations of *B. burgdorferi* fitness using the Gamma distribution to describe tick emergence instead of the stricter Uniform distribution. These simulations demonstrate that the shape of the distribution does not significantly change the qualitative results presented in the main text. We also present numerical simulations where we relax the assumption that mouse population sizes are the same across seasons. These simulations demonstrate that mouse population size does not alter the results qualitatively.

**Figure A-1:**
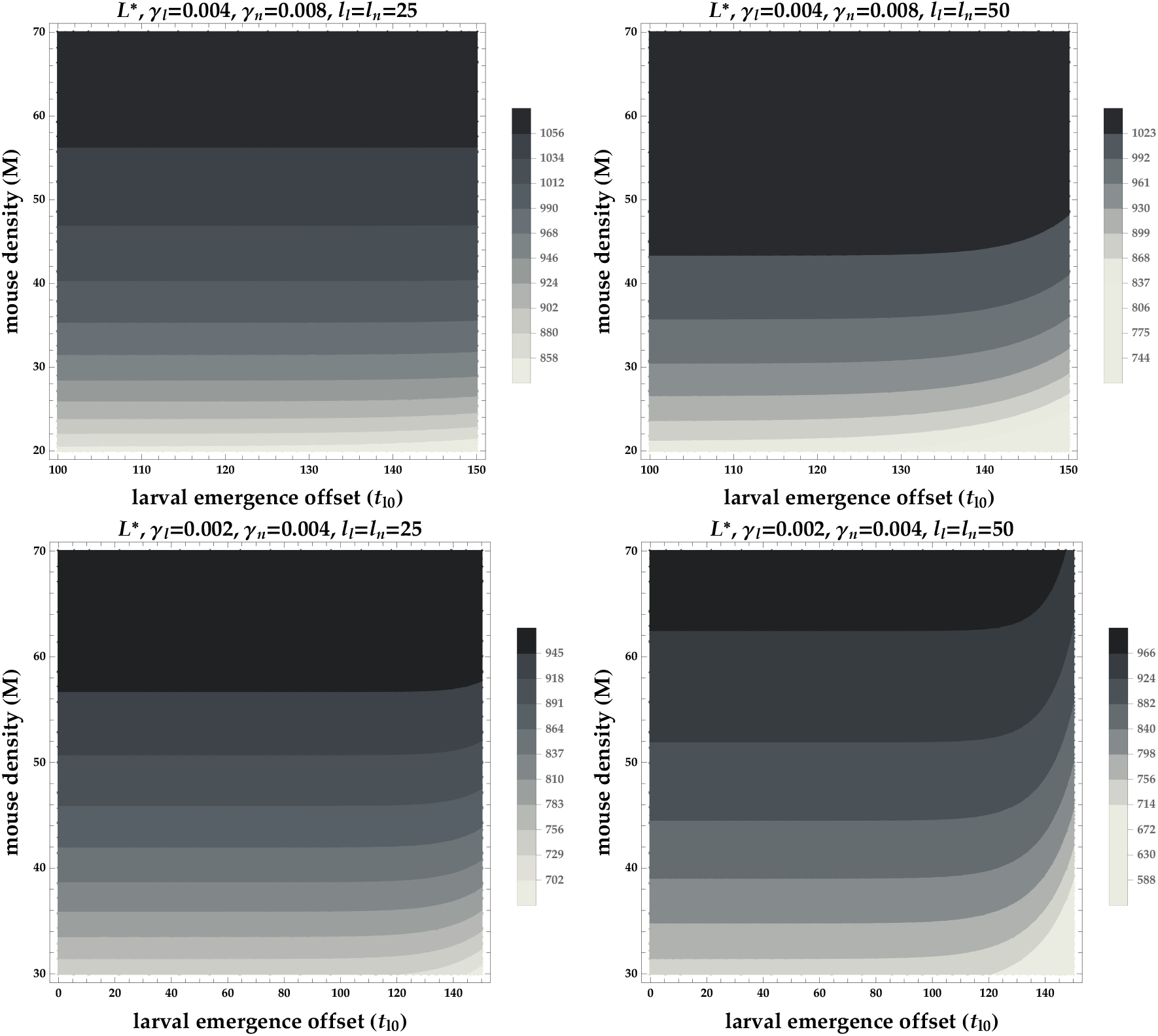
Equilibrium larval population sizes 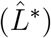 decrease at low mouse densities when larval activity begins much later than nymphal activity. *M* = *k*(1 – *μ_m_/b*) is the mouse population size, *t*_*l*0_ is the offset between when nymphs and larvae begin emerging, *γ_l_* and *γ_n_* are the contact rates between larvae and mice and nymphs and mice respectively. The first row shows that if *γ_l_* and *γ_n_* are high, large *t*_*l*0_ decreases 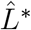 slightly when nymphal and larval emergence is broad (*l_l_* and *l_n_*). The second row shows that if *γ_l_* and *γ_n_* are low, large *t*_*l*0_ decreases 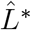 more strongly, especially when nymphal and larval emergence is broad (*l_l_* and *l_n_*). Note that contour colors are not the same across plots. All other parameter values are shown in Table 1.

**Figure A-2:**
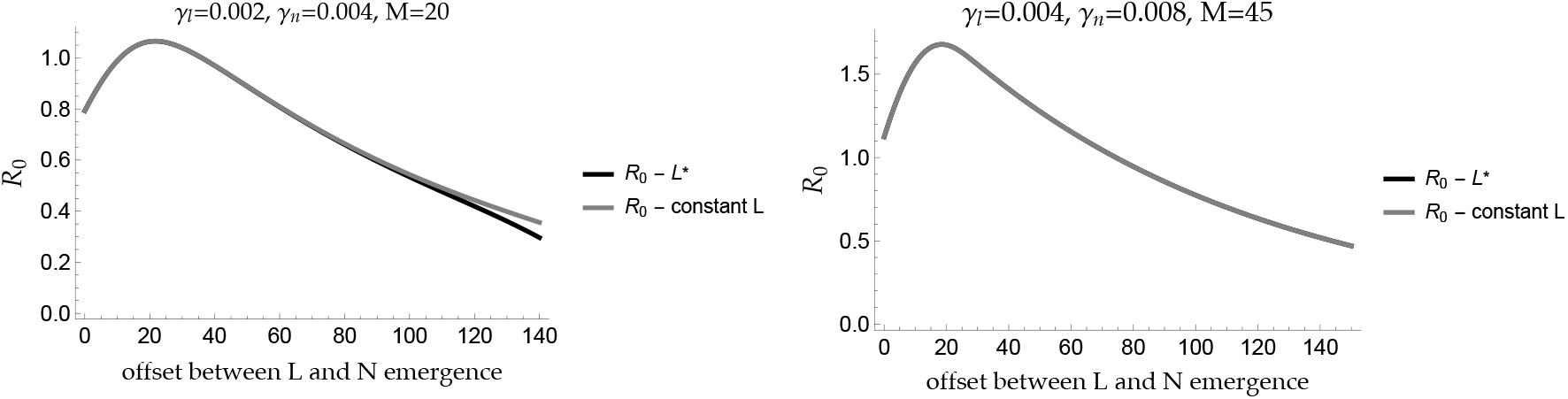
The difference between *R*_0_ that takes into account 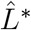 (black line) and *R*_0*nc*_ that assumes constant 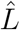 irregardless of phenology (gray line) is most dramatic for large offset between nymphal and larval activity, low contact rates between ticks and mice (*γ_l_* = 0.002, *γ_n_* = 0.004) and low mouse density (*M* = 20). The difference between *R*_0_ and *R*_0*nc*_ is negligible for higher contact rates (*γ_l_* = 0.004, *γ_n_* = 0.008) and higher mouse density (*M* = 45). 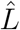 was calculated using the equation (A.3) for 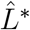 when nymphal and larval phenology is synchronous (*t*_*l*0_ = 0). As the offset between nymphal and larval activity increases, 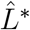 and 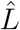 diverge driving differences in estimates for *R*_0_ and *R*_0*nc*_.

**Figure A-3:**
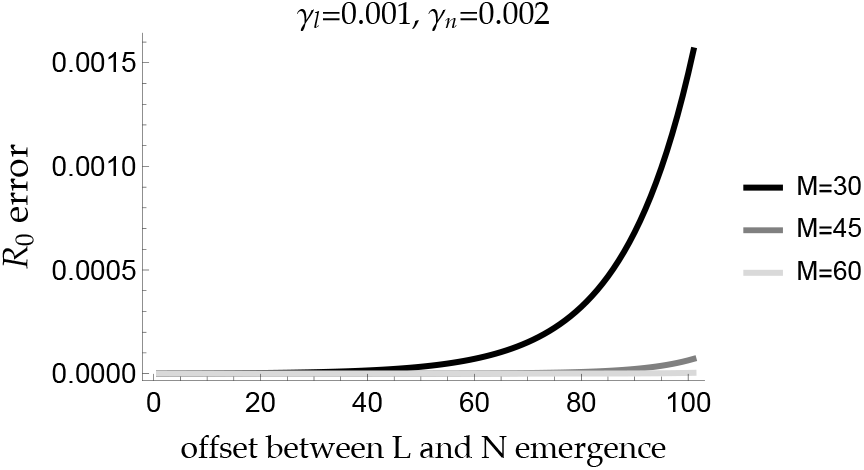
The relative error between *R*_0_ and *R*_0*nc*_ increases as the offset between larval and nymphal emergence increases. This error is exacerbated for low host density and low contact rates between ticks and mice because fewer ticks successfully feed by the end of the season which drives lower equilibrium tick sizes. *M* = *k*(1 – *μ_m_*/*b*) is the mouse population size, *γ_l_* and *γ_n_* are the contact rates between larvae and mice and nymphs and mice respectively. All other parameter values are shown in Table 1.

### Gamma distributed tick emergence

A more natural tick emergence function follows the Gamma distribution:

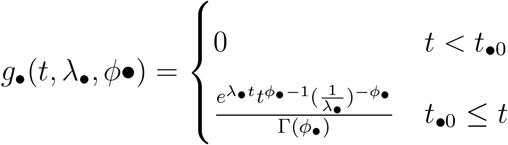

where emergence begins at *t* = *t*_•0_, *ϕ*_•_ is the shape parameter and λ_•_ is the scale parameter. We vary λ_•_ in Figures A-5 and A-6 to alter tick emergence width. Figures A-5 and A-6 show how *R*_0_ varies as a function of the time between the start of nymphal and larval emergence (*t*_*l*0_), the larval emergence width parameter (λ_*l*_) and the nymphal emergence width parameter (λ_*n*_).

Larval emergence is sometimes bimodal (Brunner and Ostfeld 2008), we thus also model larval emergence using the following distribution:

**Figure A-4:**
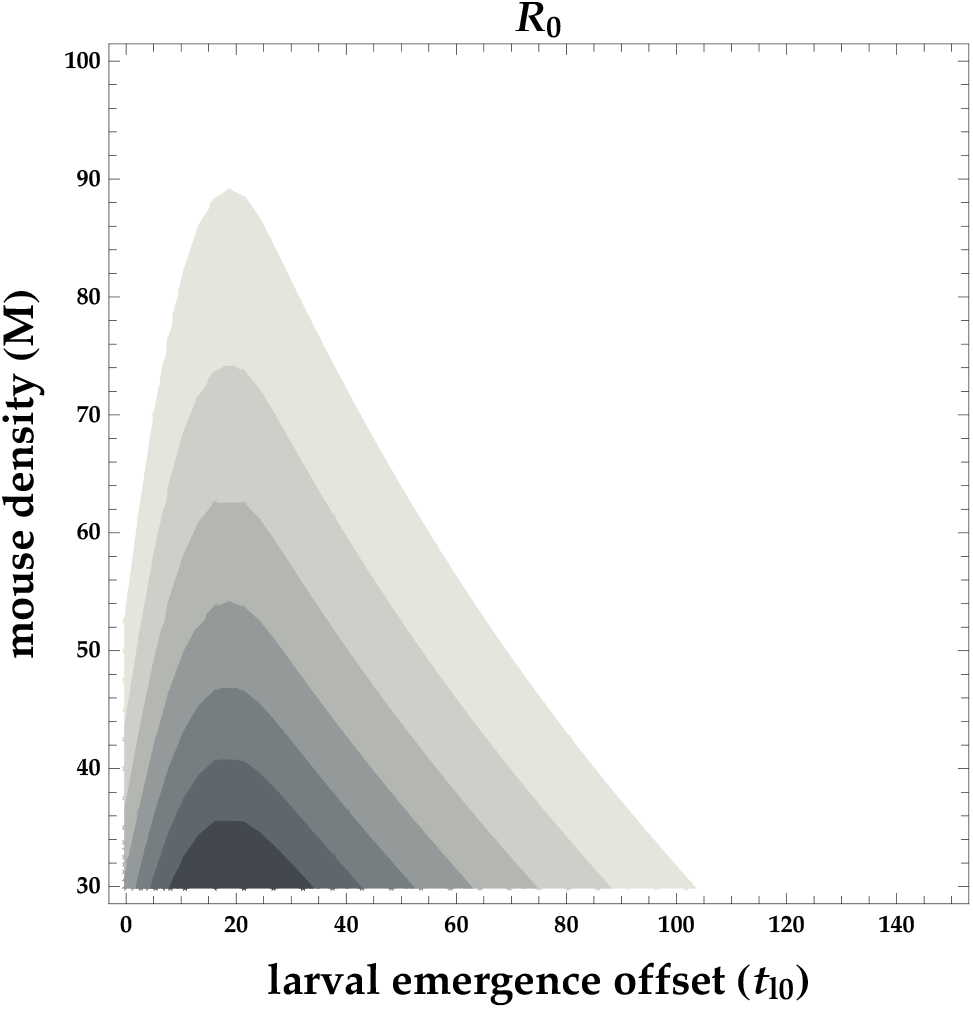
*R*_0_ decreases as mouse density increases because of dilution. *R*_0_ decreases when larval actvity begins much later than nymhpal activity because of mouse turnover.

**Figure A-5:**
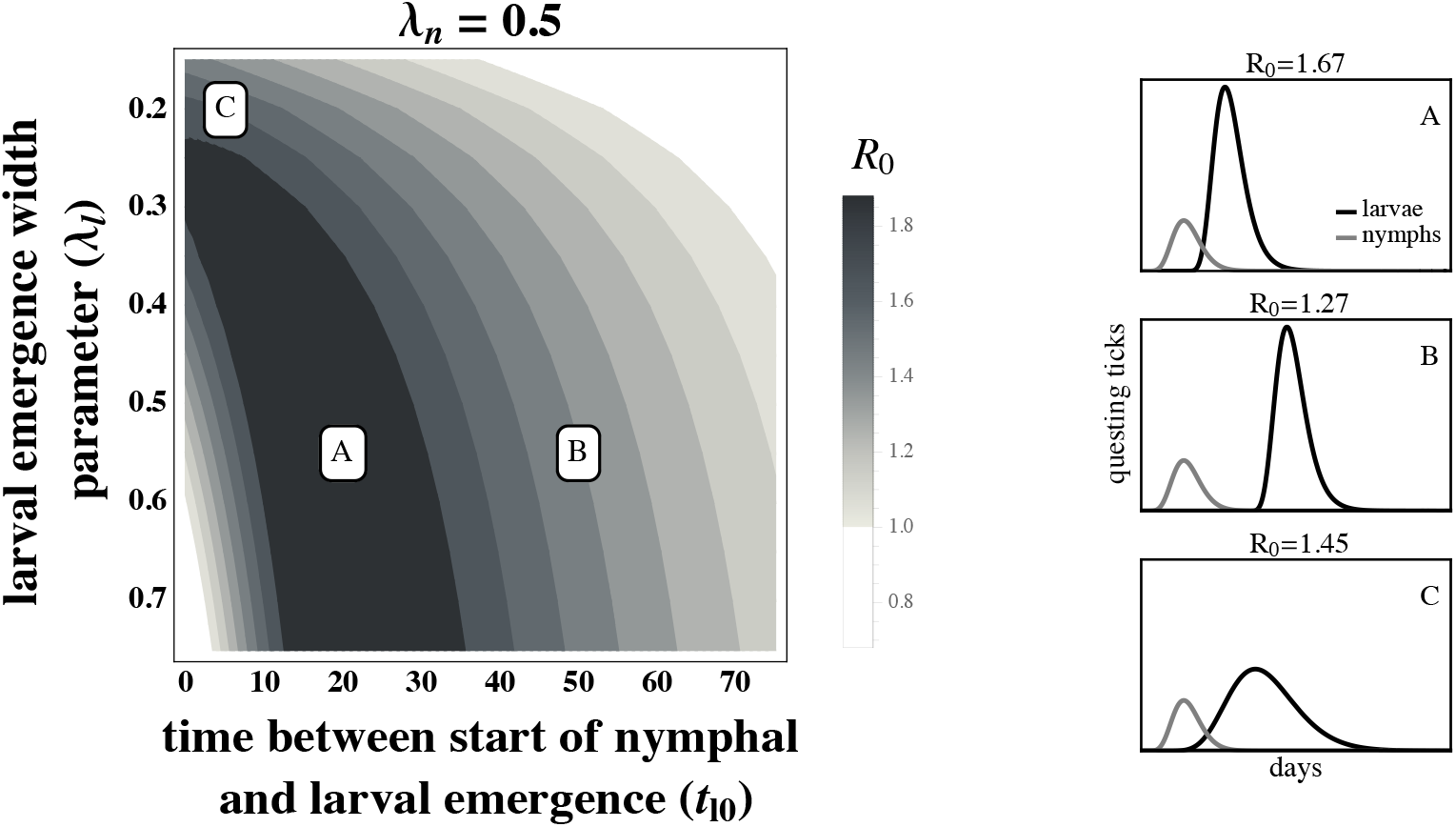
The basic reproductive number, *R*_0_, of *B. burgdorferi* is greatest when larval emergence begins shortly after nymphal emergence such that larvae are active during the peak in mouse infection prevalence. The left hand panel depicts *R*_0_ as a function of the time between the start of nymphal and larval emergence (*t*_*l*0_), and the larval emergence width parameter (λ_*l*_), and the letters indicate the parameters for the within-season dynamics on the right hand panels. (*A*) Concentrated larval emergence (large λ_*l*_) coupled with a slight difference between when nymphs and larvae begin emerging (*t*_*l*0_ < 35) increases the probability that questing larvae feed on mice recently infected by nymphs (*t*_*l*0_ = 20, λ_*l*_ = 0.55). (*B*) Greater differences between when nymphs and larvae begin emerging (*t*_*l*0_ > 35) results in lower mouse-to-larvae transmission rates as many mice infected by nymphs die and are replaced by mice born uninfected such that larvae are likely to feed on uninfected mice (*t*_*l*0_ = 50, λ_*l*_ = 0.55). (*C*) Synchronous emergence (*t*_*l*0_ = 0) can also reduce *B. burgdorferi* fitness when larval emergence duration is long (small λ_*l*_) as many larvae feed after infected mice have died (*t*_*l*0_ = 5,λ_*l*_ = 0.2). *R*_0_ is calculated assuming tick emergence is Gamma distributed. λ_*n*_ = 0.5, *ϕ_l_* = *ϕ_n_* = 10, 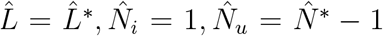 (see Appendix A). All other parameter values are shown in Table 1.

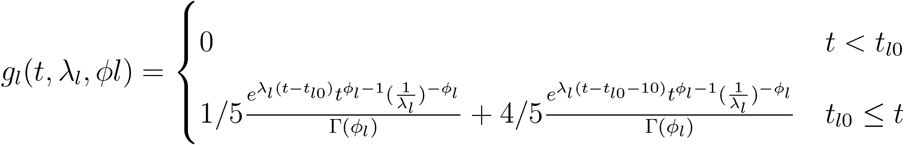

Figures A-7 and A-8 show how *R*_0_ varies as a function of the time between the start of nymphal and larval emergence (*t*_*l*0_), the larval emergence width parameter (λ_*l*_) and the nymphal emergence width parameter (λ_*n*_) when larval emergence is bimodal.

**Figure A-6:**
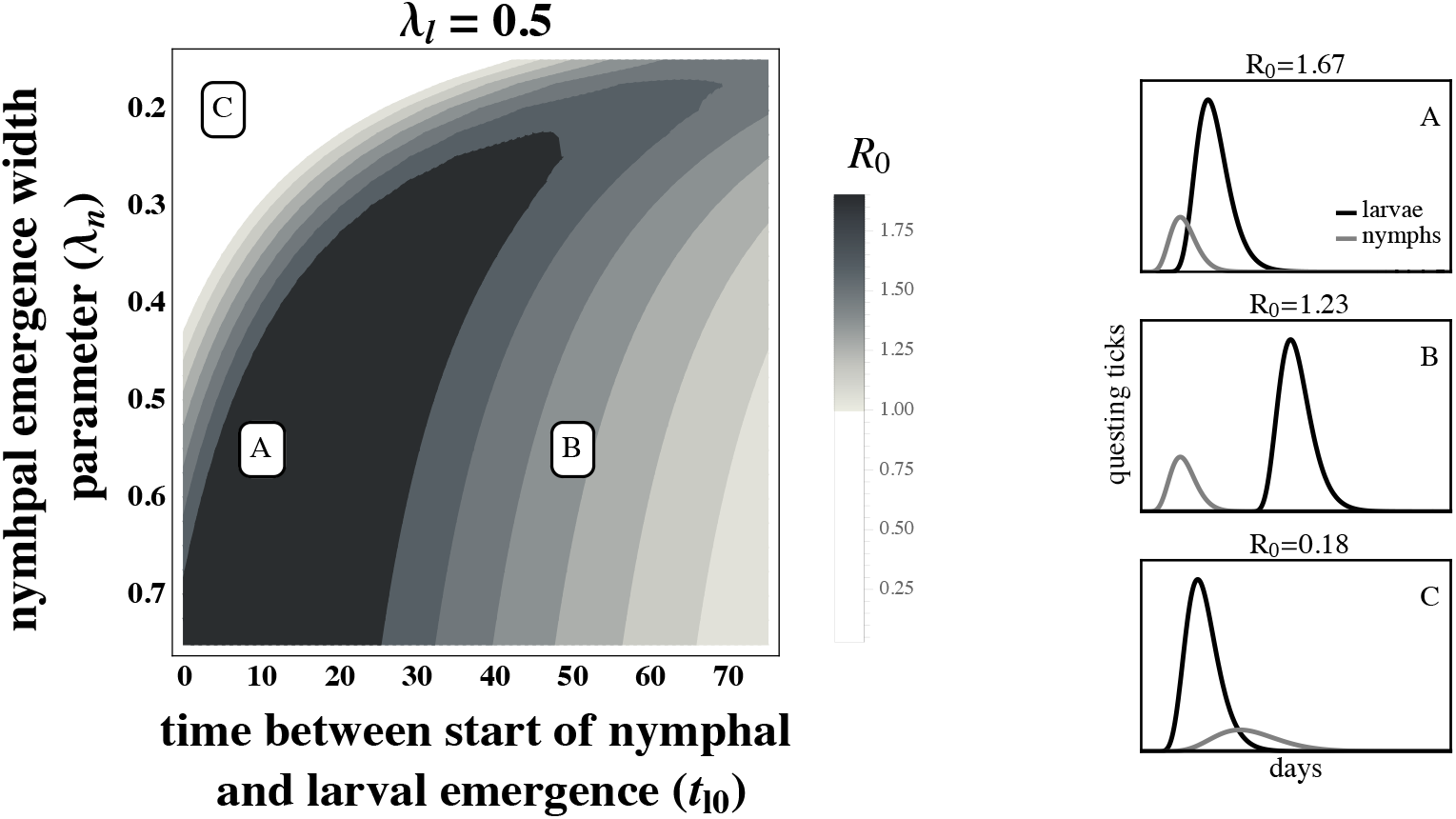
The basic reproductive number, *R*_0_, of *B. burgdorferi* is greatest when larval emergence begins shortly after nymphal emergence such that larvae are active during the peak in mouse infection prevalence. The left hand panel depicts *R*_0_ as a function of the time between the start of nymphal and larval emergence (*t*_*l*0_), and the nymphal emergence width parameter (λ_*n*_), and the letters indicate the parameters for the within-season dynamics on the right hand panels. (*A*) Concentrated nymphal emergence (large λ_*n*_) coupled with a slight difference between when nymphs and larvae begin emerging (*t*_*l*0_ < 25) increases the probability that questing larvae feed on mice recently infected by nymphs (*t*_*l*0_ = 20, λ_*n*_ = 0.55). (*B*) Greater differences between when nymphs and larvae begin emerging (*t*_*l*0_ > 25) results in lower mouse-to-larvae transmission rates as many mice infected by nymphs die and are replaced by mice born uninfected such that larvae are likely to feed on uninfected mice (*t*_*l*0_ = 50, λ_*n*_ = 0.55). (C) Synchronous emergence (*t*_*l*0_ = 0) can also reduce *B. burgdorferi* fitness when nymphal emergence duration is long (small λ_*n*_) as many larvae feed before nymphs infect mice (*t*_*l*0_ = 5, λ_*n*_ = 0.2). *R*_0_ is calculated assuming tick emergence is Gamma distributed. λ_*l*_ = 0.5, *ϕ_l_* = *ϕ_n_* = 10, 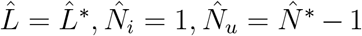 (see Appendix A). All other parameter values are shown in Table 1.

### Variable mouse density

The main mammalian host of *B. burgdorferi*, *Peromyscus leucopus*, often has variable density from year to year, (Ostfeld et al. 1996b). We relax the assumption that mouse density is constant across years by simulating how randomly varying mouse density from one year to the next impacts parasite fitness given different phenological patterns. These results are shown in Figure A-9.

**Figure A-7:**
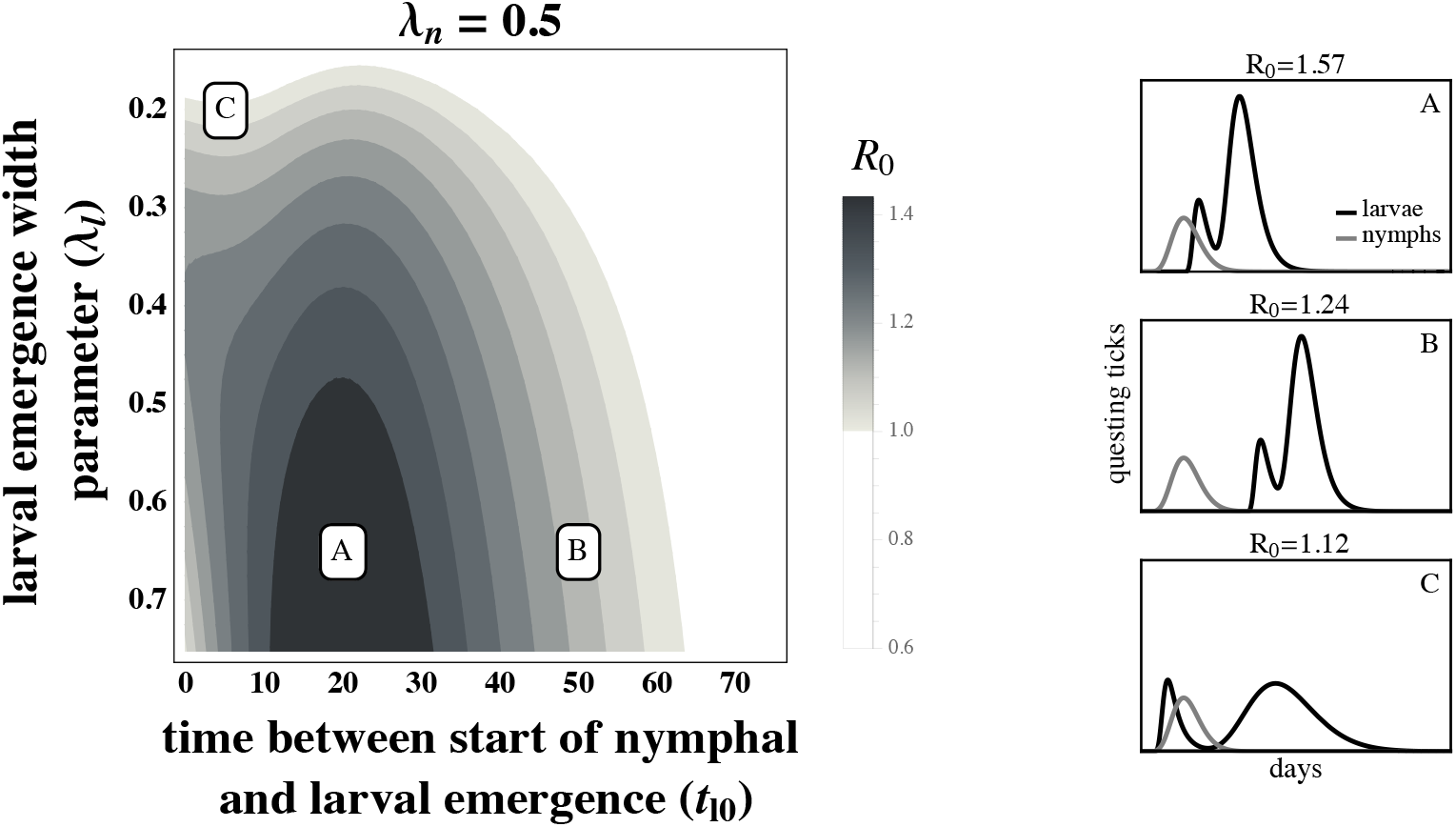
The basic reproductive number, *R*_0_, of *B. burgdorferi* is high when larvae emerge shortly after nymphs emerge so that larvae are active when mouse infection prevalence is high, despite bimodal larval emergence. The left hand panel depicts *R*_0_ as a function of the time between the start of nymphal and larval emergence (*t*_*l*0_), and the larval emergence width parameter (λ_*l*_), and the letters indicate the parameters for the within-season dynamics on the right hand panels. (*A*) Concentrated larval emergence (large λ_*l*_) coupled with a slight difference between when nymphs and larvae begin emerging (*t*_*l*0_ < 30) increases the probability that questing larvae feed on mice recently infected by nymphs (*t*_*l*0_ = 20, λ_*l*_ = 0.65). (*B*) Greater differences between when nymphs and larvae begin emerging (*t*_*l*0_ > 30) results in lower mouse-to-larvae transmission rates as many mice infected by nymphs die and are replaced by mice born uninfected such that larvae are likely to feed on uninfected mice (*t*_*l*0_ = 50, λ_*l*_ = 0.65). (*C*) Synchronous emergence (*t*_*l*0_ = 0) can also reduce *B. burgdorferi* fitness when larval emergence duration is long (small λ_*l*_) as many larvae in the first peak feed before nymphs infect mice and many larvae in the later peak feed after infected mice have died (*t*_*l*0_ = 5, λ_*l*_ = 0.2). *R*_0_ is calculated assuming nymphal emergence is Gamma distributed and larval emergence has a bimodal Gamma distribution. λ_*n*_ = 0.5, *ϕ_l_* = *ϕ_n_* = 10, 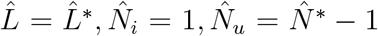 (see Appendix A). All other parameter values are shown in Table 1.

**Figure A-8:**
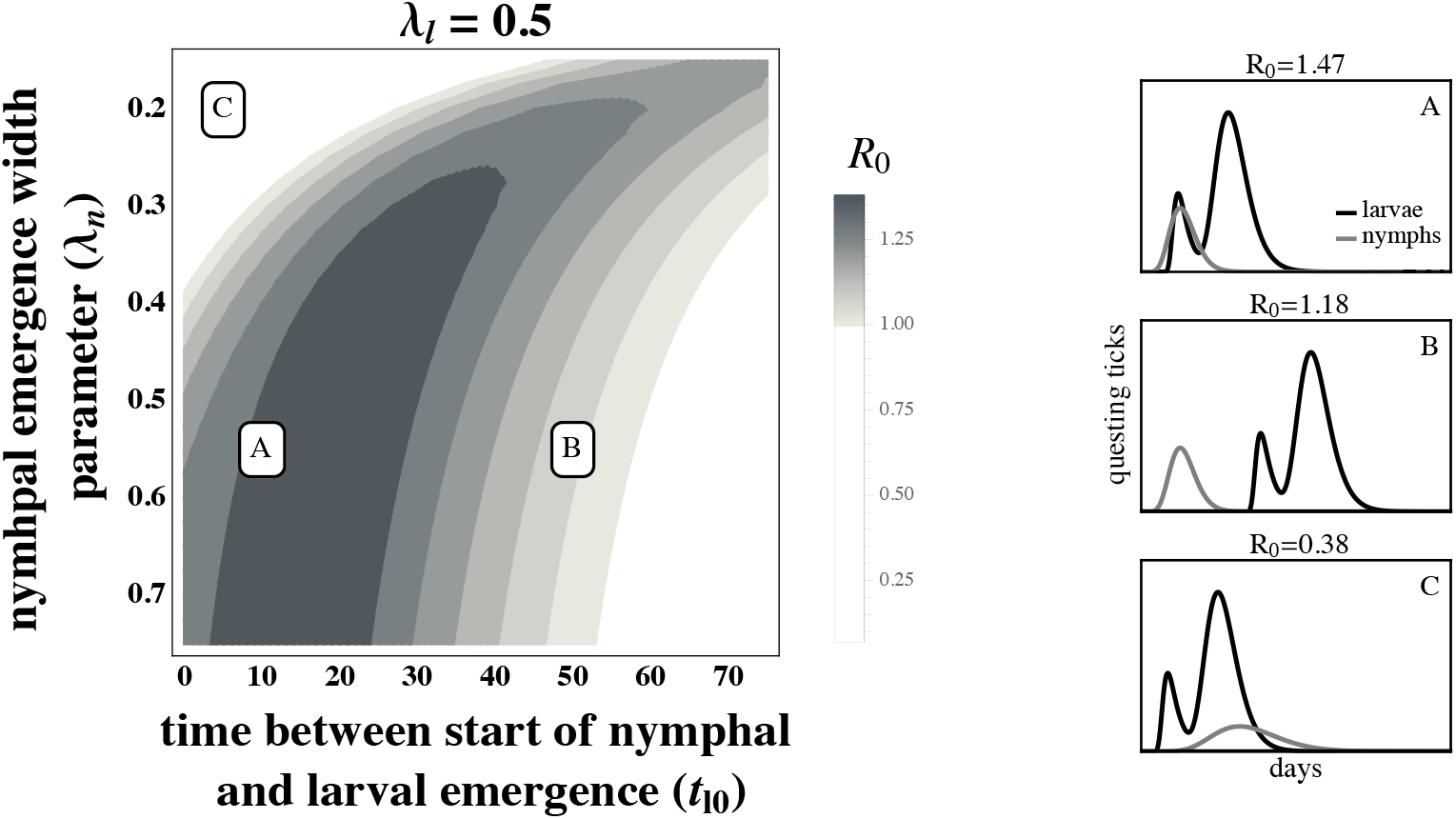
The basic reproductive number, *R*_0_, of *B. burgdorferi* is high when larvae emerge shortly after nymphs emerge so that larvae are active when mouse infection prevalence is high, despite bimodal larval emergence. The left hand panel depicts *R*_0_, in this case as a function of the time between the start of nymphal and larval emergence (*t*_*l*0_), and the duration of nymphal emergence period (λ_*n*_), and the letters indicate the parameters for the within-season dynamics on the right hand panels. (*A*) Concentrated nymphal emergence (large λ_*n*_) coupled with a slight difference between when nymphs and larvae begin emerging (*t*_*l*0_ < 25) increases the probability that questing larvae feed on mice recently infected by nymphs (*t*_*l*0_ = 10, λ_*n*_ = 0.55). (*B*) Greater differences between when nymphs and larvae begin emerging (*t*_*l*0_ > 25) results in lower mouse-to-larvae transmission rates as many mice infected by nymphs die and are replaced by mice born uninfected such that larvae are likely to feed on uninfected mice (*t*_*l*0_ = 50, λ_*n*_ = 0.55). (*C*) Synchronous emergence (*t*_*l*0_ = 0) can also reduce *B. burgdorferi* fitness when nymphal emergence duration is long (small λ_*n*_) as many larvae feed before nymphs infect mice (*t*_*l*0_ = 5, λ_*n*_ = 0.2). *R*_0_ is calculated assuming nymphal emergence is Gamma distributed and larval emergence has a bimodal Gamma distribution. λ_*l*_ = 0.5, 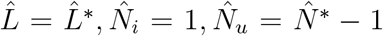 (see Appendix A). All other parameter values are shown in Table 1.

**Figure A-9:**
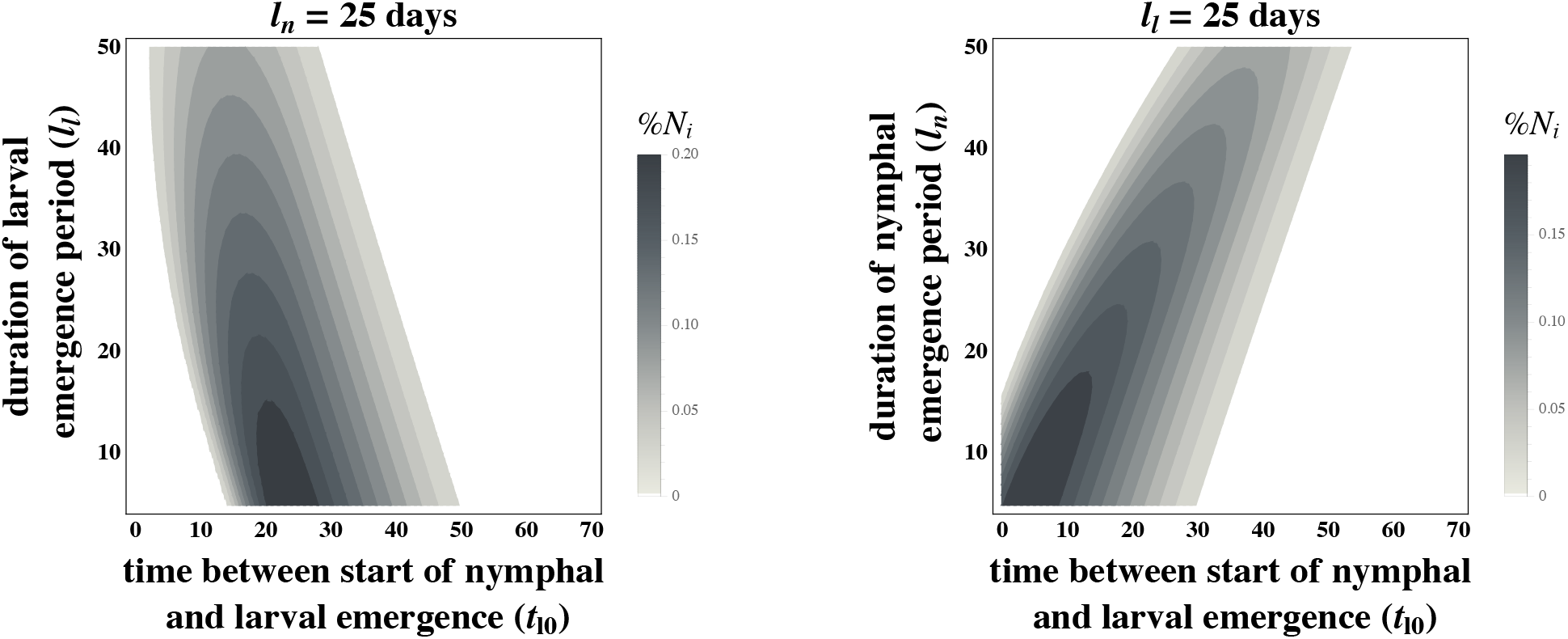
Nymphal infection prevalence (%*N_i_*) is highest when larval emergence is concentrated and begins slightly after nymhpal emergence depsite mouse density varying from one year to the next. This is the same qualitative result as shown in Figures 3 and 4 in the main text. %*N_i_* was calculated by taking the geometric mean of nymphal infection prevalence across 180 seasons when mouse density, *M*, randomly varied between 20 and 120 mice. All other parameter values are shown in Table 1.

## Notes

### Competing Interest Statement

The authors have declared no competing interest.

